# New insights into the human 26S proteasome function and regulation

**DOI:** 10.1101/2021.10.03.462214

**Authors:** Miglė Kišonaitė, Pavel Afanasyev, Jonida Tafilaku, Ana Toste Rêgo, Paula C. A. da Fonseca

**Affiliations:** MRC Laboratory of Molecular Biology, Francis Crick Avenue, Cambridge Biomedical Campus, Cambridge CB2 0QH, UK; Institute of Molecular Cell and Systems Biology, 351 Jarrett Building, Garscube Campus, University of Glasgow, Glasgow G61 1QH, UK

**Keywords:** proteasome, cryo-EM, structure, calcium, magnesium, ubiquitin, degron, regulation, function, conformation

## Abstract

The 26S proteasome is a protease complex essential for proteostasis and strict regulation of diverse critical physiological processes, the mechanisms of which are still not fully described. The human 26S proteasome purification was optimized without exogenous nucleotides, to preserve the endogenous nucleotide occupancy and conformation of its AAA-ATPase subunits. This unveiled important effects on the proteasome function and structure resulting from exposure to Ca^2+^ or Mg^2+^, with important physiological implications. This sample, with an added model degron designed to mimic the minimum canonical ubiquitin signal for proteasomal recognition, was analysed by high-resolution cryo-EM. Two proteasome conformations were resolved, with only one capable of degron binding. The structural data show that this occurs without major conformation rearrangements and allows to infer into the allosteric communication between ubiquitin degron binding and the peptidase activities. These results revise existing concepts on the 26S proteasome function and regulation, opening important opportunities for further research.

## Introduction

The 26S proteasome is a proteolytic complex essential in all eukaryotes, both for cell homeostasis and for the tight regulation of numerous and diverse fundamental cellular processes, including cell cycle progression, protein quality control, DNA repair and apoptosis. (Bhattacharyya et al., 2014; Collins and Goldberg, 2017; Livneh et al., 2016). Its canonical substrates are tagged for degradation by ubiquitin signals, typically as K48-linked tetra-ubiquitin chains, that are required for the substrate specific recognition (Thrower et al., 2000). Additionally, 26S proteasome substrates must have a disordered region required for engagement and initiation of its unfolding and translocation into the proteasome proteolytic sites (Aufderheide et al., 2015; Fishbain et al., 2015; Prakash et al., 2004; Yu et al., 2016).

The 26S proteasome is composed of at least 32 distinct canonical subunits that assemble into two major subcomplexes, the 20S proteolytic core (20S-PC) and the 19S regulatory particle (19S-RP). The eukaryotic 20S-PC is formed from two copies of α and β hetero-heptameric rings that stack into a α(1-7)β(1-7)β(1-7)α(1-7) barrel shaped assembly (Groll et al., 1997). The proteolytic active sites, associated with the subunits β1, β2 and β5, are located within the 20S-PC central inner chamber. Substrates are specifically recognized, unfolded, deubiquitinated, and translocated into this proteolytic chamber by the 19S-RP (Bhattacharyya et al., 2014; Collins and Goldberg, 2017). Biochemically, the 19S-RP can be further divided into two sub-complexes, the base and the lid (Glickman et al., 1998), the overall organization of which was first determined by cryo-electron microscopy (cryo-EM) at medium resolution (Beck et al., 2012; da Fonseca et al., 2012; Glickman et al., 1998; Lander et al., 2012). The base is composed of an unfoldase/translocase AAA-ATPase hetero-hexamer, formed by the subunits Rpt1-6, located over the adjacent 20S-PC α-ring, together with the proteasome largest subunits, Rpn1 and Rpn2. The lid comprises six peripheral PCI-motif subunits (Rpn3, Rpn5-7, Rpn9, and Rpn12) and Sem1/Rpn15, that act as scaffolding, together with the Rpn8-Rpn11 hetero-dimer. Rpn11, the constitutive deubiquitinase of the 26S proteasome, is located adjacent to the Rpt subunits where it releases the substrate ubiquitin tag by cleavage of its proximal ubiquitin (Lee et al., 2011; Verma et al., 2002). The canonical ubiquitin receptor Rpn10 is found at the periphery of the complex. Additionally, Rpn1 and Rpn2 can dock various proteasome adaptors including shuttling factors (Rad23, Dsk10), additional ubiquitin receptors (Rpn13) and ancillary deubiquitinases (Ubp6, Usp14) that release substrate degron distal ubiquitin moieties (Lee et al., 2011; Shi et al., 2016). It was also reported that Rpn1 can also directly act as an ubiquitin receptor (Shi et al., 2016). High resolution cryo-EM structures of complete apo 26S proteasomes were reported, including at resolutions higher than 4 Å (Chen et al., 2016; Huang et al., 2016; Zhu et al., 2018). Additionally, 26S proteasome structures were also determined in the presence of added substrates, where degradation was stalled either by using non-hydrolysable nucleotide analogues or by introducing proteasome mutations (de la Peña et al., 2018; Ding et al., 2019; Dong et al., 2019). These studies resolved distinct proteasome conformations that have been interpreted mainly in terms of a model describing substrate engagement, ATP-hydrolysis steps and their mechanochemical coupling to substrate unfolding, leaving the detailed mechanisms of how the 26S recognizes ubiquitinated substrates still elusive.

Early studies showed that the presence of ATP, but not its hydrolysis, is required for the *in vitro* preservation of stable 26S proteasomes (Eytan et al., 1989; Ganoth et al., 1988; Hough et al., 1987; Liu et al., 2006), which otherwise dissociate into their 20S-PC and 19S-RP components. Therefore, the purification of 26S proteasomes is generally, if not always, performed with ATP (or an analogue) and MgCl_2_ added to all solutions. However, the use of an excess of exogenous nucleotides may interfere with the endogenous occupancy of the ATPase active sites, with consequences to the overall conformation of the complex. Here we present the characterization of the human 26S proteasome prepared in the absence of exogenous nucleotides. Remarkably, the optimization of the 26S proteasome purification protocol unveiled an unexpected modulation of the 26S proteasome peptidase activities and structural integrity by Ca^2+^ and Mg^2+^, with important physiological implications. The analysis of this sample in the presence of a minimal ubiquitin-based model degron, optimized to form stable interactions with the proteasome, provided insights into the proteasome transient ubiquitin-degron recognition step that precedes substrate engagement and commitment to degradation.

## Results

### Preparation of human 26S proteasomes in the absence of exogenous nucleotides

The preparation of human 26S proteasomes was optimized in the absence of exogenous nucleotides, resulting in the protocol described in the Methods section. Initially, it was observed that just removing the nucleotides from the solutions resulted in the dissociation of the 26S proteasome, as previously reported (Eytan et al., 1989; Ganoth et al., 1988; Hough et al., 1987). However, during the protocol optimization it was observed that it was also necessary to exclude MgCl_2_ and add EDTA to all purification solutions, particularly during cell lysis, in order to obtain stable complexes in the absence of exogenous nucleotides. By using this protocol, we obtained 26S proteasomes with a protein composition indistinguishable from conventionally prepared proteasomes, as judged by SDS-PAGE (Figure 1A) and mass-spectrometry. These complexes are active for both unfolding and proteolysis, as shown by the degradation of a previously reported model folded protein substrate (Bhattacharyya et al., 2016) (Figure 1B). However, the individual peptidase activities of 26S proteasomes purified without exogenous nucleotides differed from those of conventional samples, as described below.

**Figure 1.**
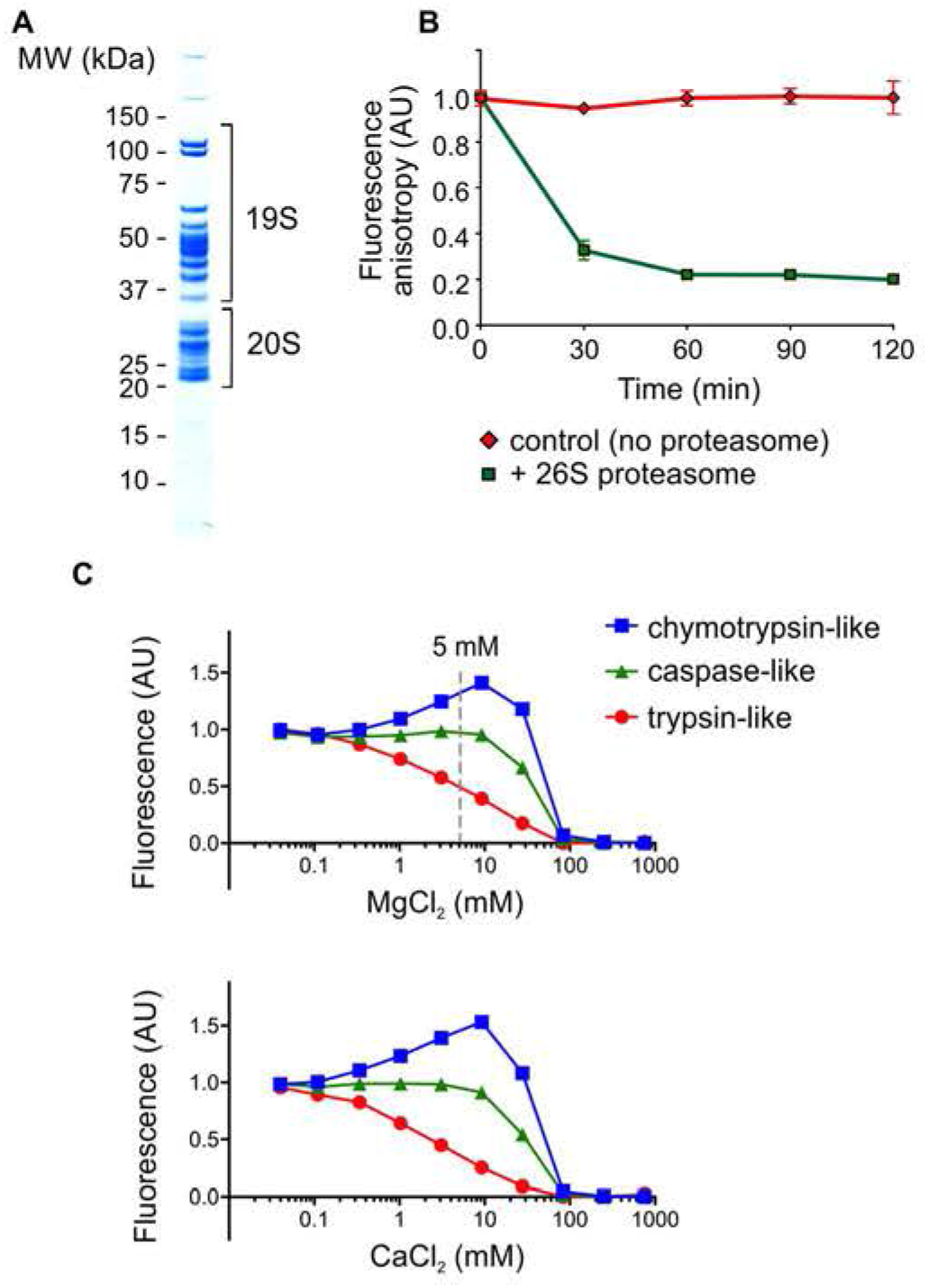
26S proteasome sample characterization and the effects of Ca^2+^/Mg^2+^ on its peptidase activity. **(A)** SDS-PAGE of the optimized 26S proteasome sample prepared in the absence of exogenous nucleotides. **(B)** 26S proteasome proteolytic activity against a fluorescently labeled model folded protein substrate, incubated in the absence (red) or presence of 50 nM 26S proteasome purified without exogenous nucleotides (green). Error bars represent the standard deviation from the average of all measurements. **(C)** Effects of MgCl_2_ and CaCl_2_ on the three peptidase activities of the 26S proteasome. Each titration curve was normalized to start at a fluorescence intensity of 1 and have a minimum of 0. The concentration of 5 mM MgCl_2_, corresponding to the typical working concentration used during conventional 26S proteasome purifications in the presence of ATP (or analogues) is indicated by a dashed line.

### 26S proteasome modulation by Ca^2+^ and Mg^2+^

The optimization of the preparation of human 26S proteasomes in the absence of exogenous nucleotides clearly showed that, to obtain stable complexes, MgCl_2_ also needed to be excluded from all solutions used. These conditions are ideal to investigate the effects of Mg^2+^, and also Ca^2+^, on the proteasome function and structural integrity. When each of the proteasome peptidase activities is assessed individually using fluorogenic peptides, it is clear they are all affected by the presence of these cations (Figure 1C). Ca^2+^ and Mg^2+^ have an inhibitory effect on the proteasome caspase-like and trypsin-like activities, associated with the β1 and β2 active sites, respectively, with the trypsin-like activity reduced even at low cation concentrations. On the other hand, the chymotrypsin-like activity, associated with the β5 active site, shows a biphasic behavior with activation up to 10 mM MgCl_2_ or CaCl_2_ and inhibition at higher concentrations. These results imply that the peptidase activities of 26S proteasomes prepared in the absence of divalent cations differ from those of conventionally prepared complexes, usually maintained in the presence of 5-10 mM of MgCl_2_ (Figure 1C). Additionally, the effects of Ca^2+^ or Mg^2+^ on the structural integrity of the 26S proteasome were assessed by electron microscopy of negative stained samples, that showed a noticeable dissociation of the 26S proteasome into its 20S-PC and 19S-RP sub-complexes when exposed to increasing concentrations of CaCl_2_ or MgCl_2_ (Figure S1).

### Identification of a minimal proteasome substrate ubiquitin degron mimetic

The preparation of well-defined homogeneous poly-ubiquitin chains, either enzymatically or by chemical synthesis, is still not straightforward. To overcome this, we developed a ubiquitin-derived construct that mimics the biochemical properties of the interaction between the 26S proteasome and endogenous K48 linked tetra-ubiquitin chains, the canonical degron for proteasome recognition (Thrower et al., 2000), that can be easily and consistently prepared by recombinant expression. A series of constructs, originally based on the ubiquitin mutant UBB^+1^ (Shabek et al., 2009), was designed to comprise one or two ubiquitin moieties flanked by disordered regions of varied lengths (Figure S2). The interaction of these proteins with the 26S proteasome was evaluated by peptidase activity assays, following earlier studies that showed that the proteasome chymotrypsin-like activity is enhanced in the presence of ubiquitinated substrates (Bech-Otschir et al., 2009; Nathan et al., 2013). However, adding the ubiquitin-based constructs to the 26S proteasome, prepared without exposure to Ca^2+^ or Mg^2+^, had no effect on its chymotrypsin-like activity, even in control experiments using previously characterized substrates. On the other hand, if MgCl_2_ was added to the activity assay buffer, then peptidase activity enhancement was observed for some of the constructs tested, indicating their direct interaction with the proteasome (Figure 2A). Such activation did not occur in control assays where the constructs were added to the 20S-PC sub-complex (Figure 2B), indicating that the effects observed are not due to a non-specific direct interaction with the proteolytic core but requires the presence of 19S-RPs, where the 26S proteasome ubiquitin receptors are located. To obtain a signal indicative of degron interaction with the 26S proteasome, the activity measurements described below were therefore performed with MgCl_2_ added only to the assay solution, and not to any other stage of the proteasome purification.

**Figure 2.**
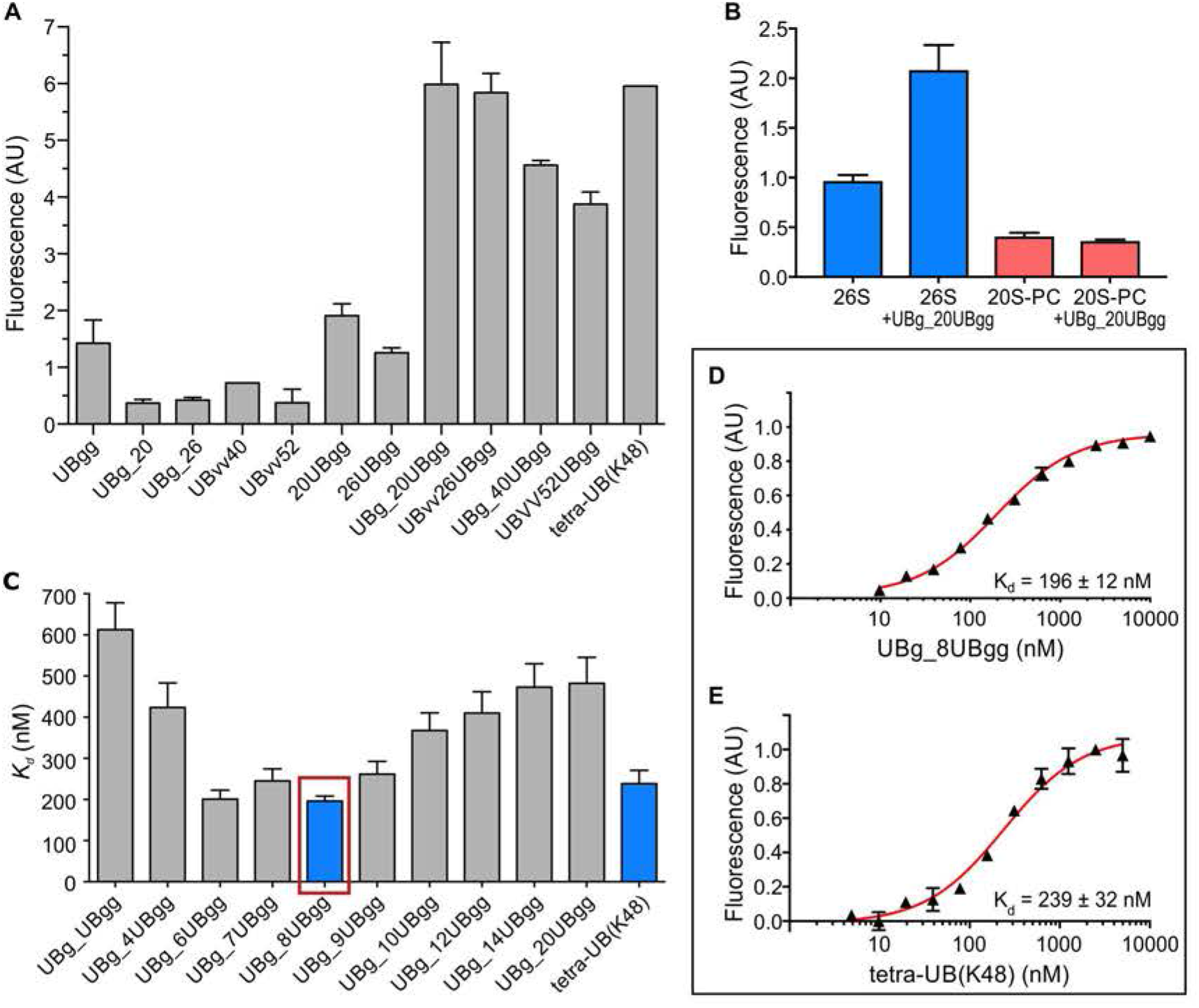
26S proteasome interaction with model ubiquitin degrons. **(A)** Chymotrypsin-like activity assays to preliminary assess the 26S proteasome interaction with the model degrons indicated, where a 26S proteasome-degron interaction results in increased fluorescence. **(B)** Control experiment, where the chymotrypsin-like activities of 20S-PC (red) and 26S proteasomes (blue) were assayed in the absence or presence of UBg_20UBgg, which interacts with the26S proteasome according to (**A**). **(C)** Binding affinities of model degrons to the 26S proteasome, used to identify an optimal ubiquitin degron mimetic (boxed in red). **(D, E)** Titration plots of the 26S proteasome chymotrypsin-like activity in the presence of increasing concentrations of UBg_8UBgg (**D**) or K48 linked tetra-ubiquitin chains (**E**), used to determine their proteasome binding affinities (see also Figures S2 and S3 and Table S1). Error bars represent the standard deviation from the average of all measurements.

The 26S proteasome chymotrypsin-like peptidase activation by the ubiquitin-derived constructs was first qualitatively tested by tracking the fluorescence intensity resulting from the cleavage of a commonly used small fluorogenic peptide (Suc-LLVY-AMC). These preliminary assays showed higher activity when constructs containing two ubiquitin moieties linked by a loop were tested, with the stronger effect observed for a construct with the smallest loop of 20 amino acids (Figure 2A). To further characterize the effects of the intervening loop on the proteasome activation, constructs with loops varying from 0 to 20 amino acid residues were tested, and for these the peptidase activity assays were used to quantitatively measure their binding affinity to the 26S proteasome (Figures 2C, S2 and S3, and Table S1). The ubiquitin-derived constructs with intervening loops of 6 to 9 amino acid residues have the highest proteasome binding affinity. Amongst these, we selected the construct with a loop formed by 8 residues (UBg_8UBgg), with a *K*_*d*_ of 196 nM. This is of the same magnitude as the *K*_*d*_ of 239 nM measured for the physiological K48 linked tetra-ubiquitin (Figure 2D and E, and Table S1), a value that is in agreement with previously published data (Thrower et al., 2000). The substitution of the UBg_8UBgg loop (residues ^76^YADLREDP^83^) by 8 alanine residues did not significantly affect the binding to the proteasome, as shown by the resulting *K*_d_ of 261 nM (construct UBg_8(polyA)UBgg, Table S1). This indicates that the loop linking the two ubiquitin moieties is not involved in major interactions with the proteasome, suggesting instead that it plays a role as a flexible spacer between the two proteasome interacting ubiquitin moieties, with its 8 residues representing the optimal length for proteasomal interaction.

### Cryo-EM of the 26S proteasome in the presence of UBg_8UBgg reveals two conformational states

Single particle analysis was used to directly infer into the interaction between the ubiquitin degron mimetic UBg_8UBgg and the 26S proteasome, purified in the absence of exogenous nucleotides and without exposure to Ca^2+^ or Mg^2+^ (Figures 1A, S4A and S5). During data processing, three-dimensional classification clearly showed that the 26S proteasome in the proteasome-UBg_8UBgg sample predominantly adopts two distinct conformations, which co-exist in approximately equal amounts (Figures 3 and S5). Broadly, these two conformations, resolved at ∼3.6 Å resolution, can be distinguished by their degron occupancy, an overall rotation of the 19S-RP, and the conformational stability of Rpn1 and Rpn10, as detailed below.

**Figure 3.**
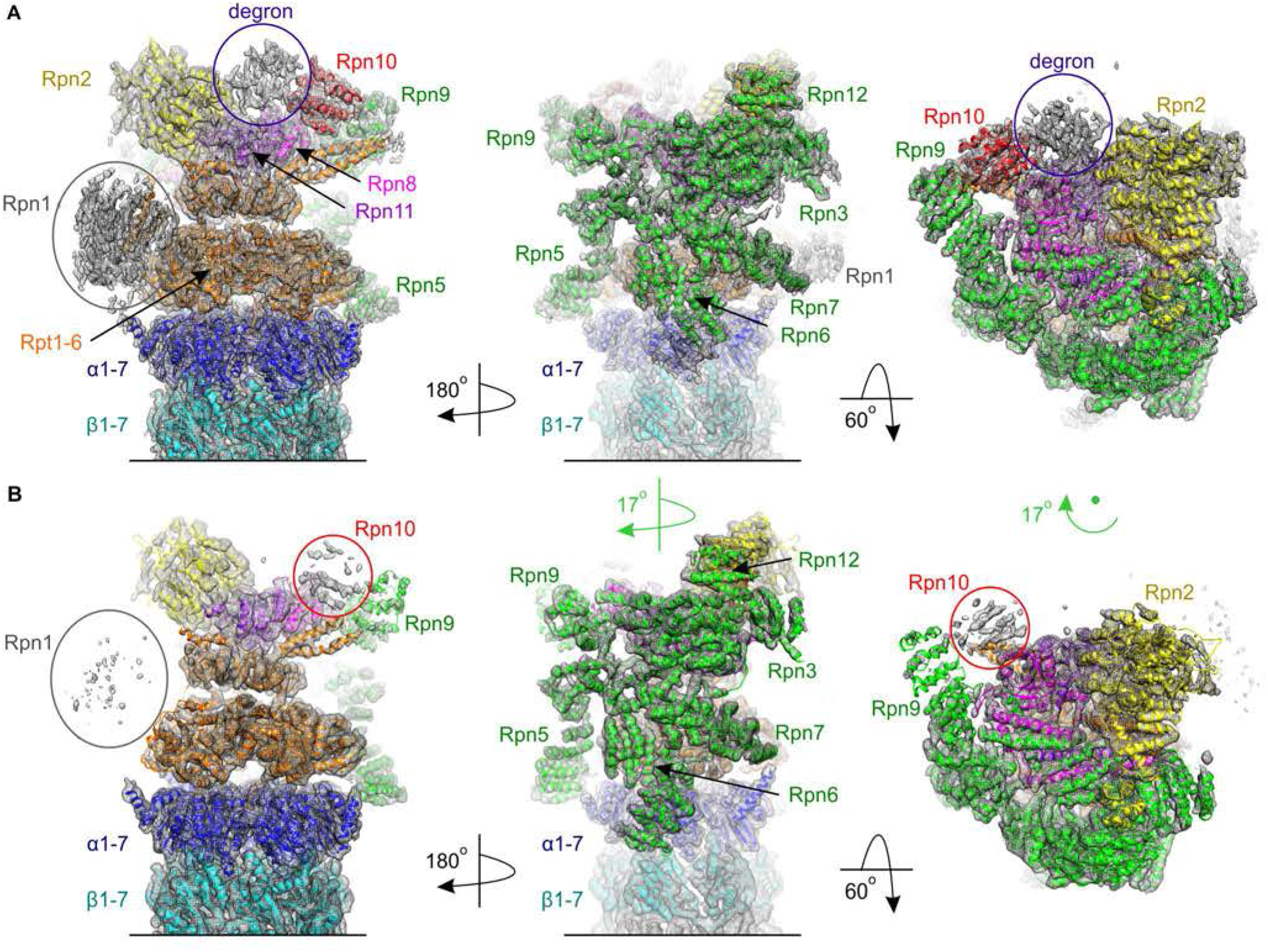
Cryo-EM structures of the human degron-binding and degron-free 26S proteasome conformations. **(A)** Cryo-EM map of the 26S proteasome degron-binding state (mesh) with fitted atomic model (cartoon representations). **(B)** Cryo-EM structure of the 26S proteasome degron-free state, represented in the same way as the data in (**A**). Different 26S proteasome subunits are indicated. The 19S-RP rotation observed between the two conformations is represented in green (see also Video S1).

When the two proteasome conformations are compared, a density bridging Rpn2 and Rpn10 (the 26S proteasome canonical ubiquitin receptor), and adjacent to Rpn11 (the proteasome constitutive deubiquitinase) is only found in one of the conformations and can be assigned to the degron UBg_8UBgg (Figure 3A). Overall, the degron-binding conformation of the 26S proteasome differs from the degron-free state by a global 17° clockwise rotation of the 19S-RP about the long axis of the 26S proteasome, as viewed towards the 20S-PC (Video S1). In the degron-binding conformation, densities for both Rpn1 and the Rpn10 N-terminal VWA domain are also significantly better recovered (Figure 3). Moreover, a closer comparison shows other noticeable differences, particularly at the Rpt hetero-hexamer. In both proteasome conformations the Rpt AAA-ATPase domains are co-planar, with their arrangement departing from a closed ring by the outward displacement of two neighboring Rpt subunits, resulting in the opening of an adjacent cleft (Figure 4). The subunits displaced outwards are Rpt5-Rpt1 in the degron-binding conformation (Figure 4A), resulting in a cleft between Rpt1 and Rpt2, while in the degron-free conformation the subunits displaced outwards are Rpt1-Rpt2, with the cleft formed between Rpt2 and Rpt6 (Figure 4B). Such arrangements are accompanied by different nucleotide occupancies (Figures 4 and S6). Despite the absence of exogenous nucleotides during the 26S proteasome purification and imaging, in the two conformations endogenous nucleotides are found at four of the AAA-ATPase active sites. In both structures the two Rpt subunits that are outwards displaced are found in the apo state (Figures 4C and S6). In the degron-binding conformation, densities compatible with ATP molecules are found at the remaining Rpt subunits, namely Rpt2-Rpt6-Rpt3-Rpt4. On the other hand, in the degron-free conformation clear nucleotide densities that can be assigned to ATP are found only at subunits Rpt6-Rpt3, with densities more consistent with ADP found at the subunits Rpt4-Rpt5, with the weaker appearance of the latter densities suggestive of some level of partial occupancy for ADP.

**Figure 4.**
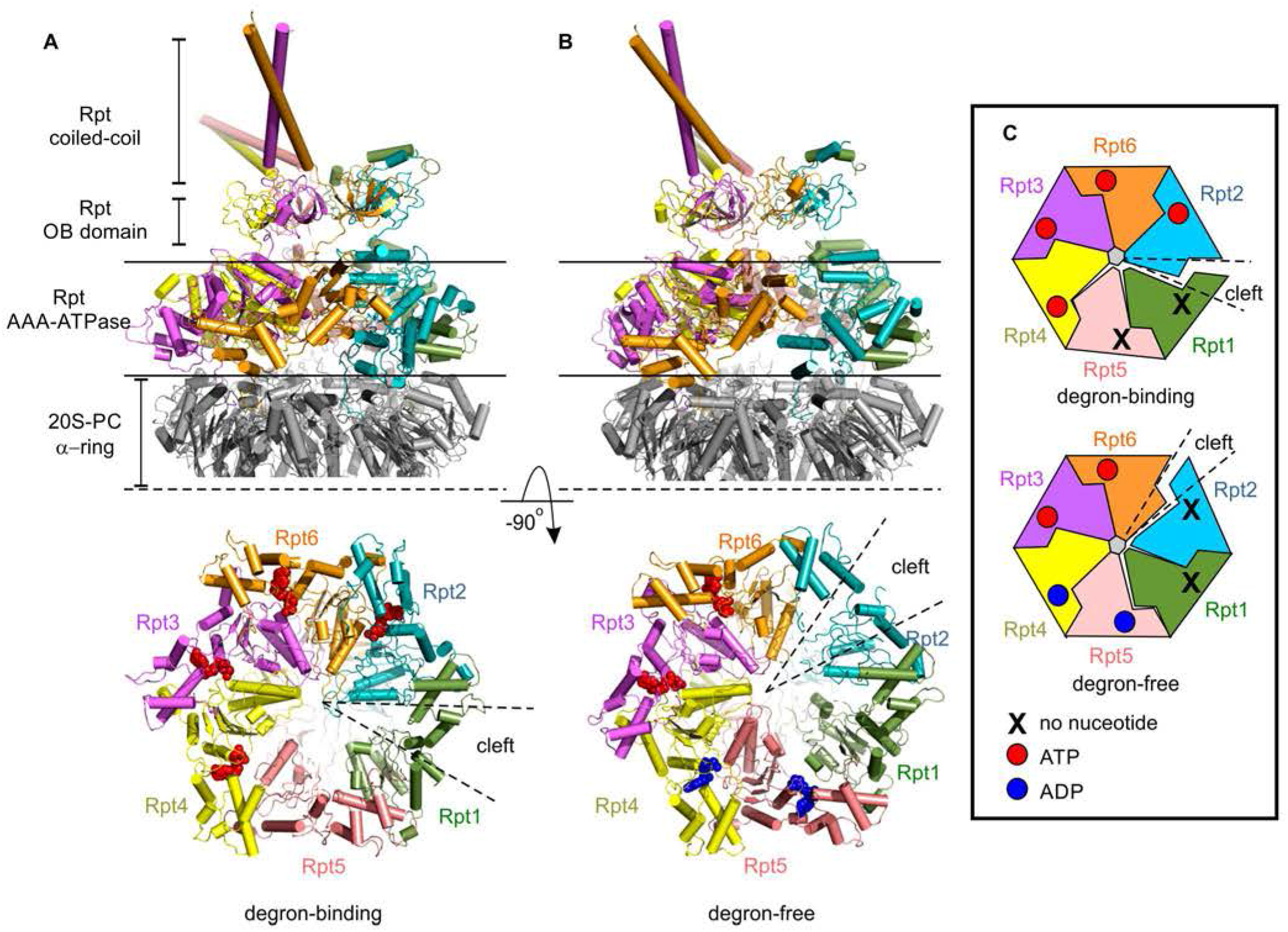
The Rpt subunit organization within the two human 26S proteasome conformations. **(A)** The Rpt subunits in the degron-binding 26S proteasome state. Top panel: side view of the protein model of the Rpt subunits and adjacent 20S-PC α-ring. Lower panel: section of the model at the Rpt subunit ring, viewed from the 20S-PC, showing the departure from a closed hetero-hexameric ring. **(B)** The Rpt subunits in the degron-free 26S proteasome state, shown as in **(A). (C)** Schematic representation of Rpt subunit arrangements and nucleotide occupancy in the human 26S proteasome degron-binding (top panel) and degron-free (lower panel) states.

Densities for each of the proteasome anchoring Rpt C-terminal HbYX (hydrophobic-tyrosine-other) motifs are recovered at their 20S-PC α-subunit ring binding pockets, except for Rpt4 for which the C-terminal tails are disordered in both conformations (Figure 5). The anchoring of the Rpt HbYX motifs at the interface of the 20S-PC α-subunits has been previously suggested to be associated with the displacement of the N-terminal loops of the α-subunits resulting in the opening of an axial channel at the 20S-PC (Smith et al., 2007). However, in the current analysis the N-terminal loops of the α-subunits are found in an open conformation in the degron-binding and closed in the degron-free state, (Figure 5), despite the similar engagement of Rpt HbYX motifs at the 20S-PC α-rings in both conformations. This suggests that the main driver for the 20S-PC channel opening is the overall conformation of the Rpt hetero-hexamer rather than the differential binding of the Rpt HbYX on the 20S-PC surface, which may otherwise be associated with the structural stability of the complex.

**Figure 5.**
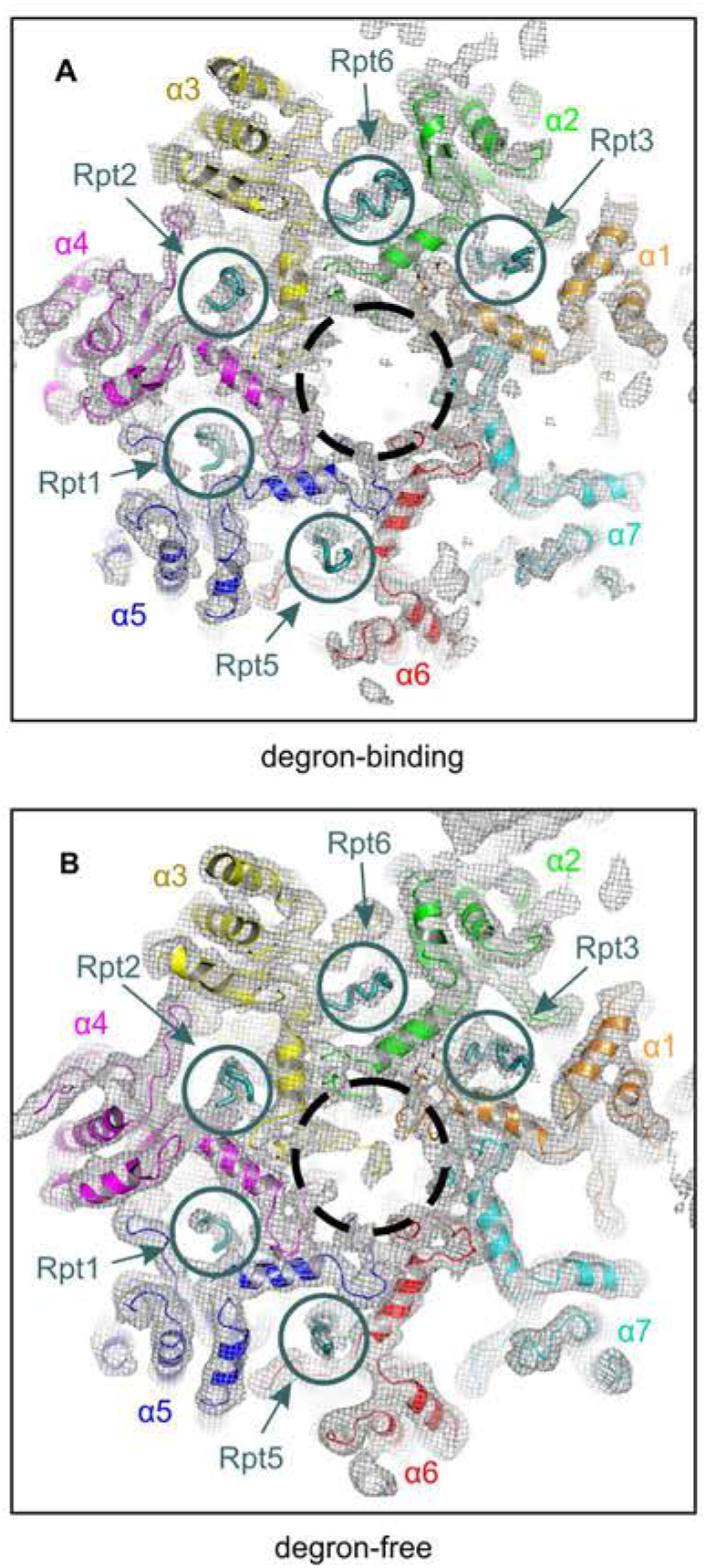
The 20S-CP α ring in the two human 26S proteasome conformations. **(A)** The 20S-PC α-ring of the human 26S proteasome degron-binding state (protein model shown as cartoon), viewed from its outer surface and along the proteasome long axis, showing the axial channel (dashed black circle) and the location of the 5 docked Rpt C-terminal HbYX motifs (solid circles, protein model shown as sticks). Each of the Rpts is named, indicated by arrows **(B)** The 20S-PC α-ring of the human 26S proteasome degron-free state, represented as in (**A**).

### The two 26S proteasome conformations in the presence of UBg_8UBgg remain in the absence of degron

The distinct features of the degron-binding and degron-free 26S proteasome conformations described here may appear to suggest a global conformational rearrangement associated with the binding of UBg_8UBgg. However, this is not supported by the distinct details of the two proteasome conformations, particularly by their distinct nucleotide binding sites occupancy as explained in the Discussion section below. Therefore, a control cryo-EM analysis of the apo 26S proteasome, in the absence of UBg_8UBgg and exogenous nucleotides, was performed to determine the consequences of the degron binding in the proteasome conformation. Apo 26S proteasomes were prepared and analyzed by single particle cryo-EM following the procedures similar to those used for the analysis of the proteasome-UBg_8UBgg sample. The two proteasome conformations resolved from the proteasome-UBg_8UBgg sample also dominate in the sample in the absence of degron, at a comparable stoichiometry and with the same nucleotide occupancies, although both lacking the UBg_8UBgg densities (Figure 6). In the apo 26S proteasome map in the degron-binding conformation, some weak densities remaining in the region so of the degron binding site can be assigned to the unboud C-terminal ubiquitin interacting motifs (UIM) of Rpn10 (Young et al., 1998), which are highly flexible in the absence of a ubiquitin degron bound (Figure 6A). These results suggest that rather than representing a conformational rearrangement associated with degron binding, the two proteasome states pre-exist with only one capable of binding UBg_8UBgg, which occurs without major proteasome rearrangements.

**Figure 6.**
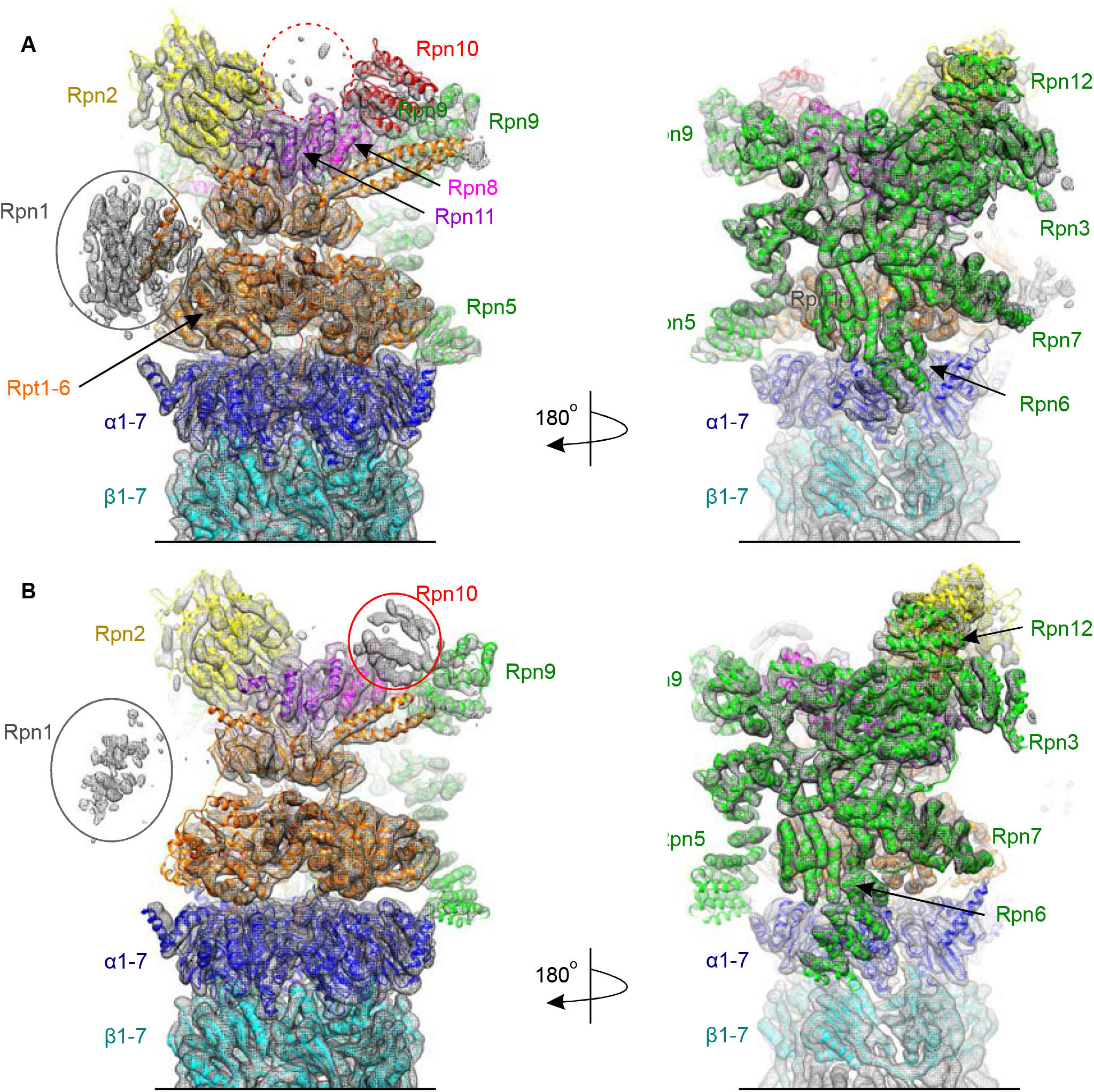
Control cryo-EM maps of apo human 26S proteasomes purified in the absence of exogenous nucleotides. Views of the cryo-EM maps (mesh representation) of the degron-binding (**A**) and degron-free (**B**) states of the human 26S proteasome, in the absence of exogenous nucleotides and ubiquitin degrons. The corresponding protein models (carton representations) derived from the cryo-EM maps obtained in the presence of UBg_8UBgg (Figure 3) were rigid-body fitted into the apo proteasome cryo-EM maps shown here. The close fit between the cryo-EM maps of the apo 26S proteasome and the protein models of the complex in the presence of degron demonstrates that no significant conformational rearrangements occur upon ubiquitin-degron binding. A dashed red circle in (**A**) indicates the location of the unoccupied ubiquitin degron binding site.

## Discussion

### 26S proteasome regulation by Ca^2+^/Mg^2+^

The characterization of the 26S proteasome functional mechanisms and regulation, including the different sequential steps of specific substrate recognition, unfolding and translocation into the 20S-PC proteolytic chamber, requires the preparation of complexes close to their endogenous state. Previously established 26S proteasome purification protocols require the presence of ATP (or a related nucleotide analogue) and MgCl_2_ in all buffer solutions. However, the presence of an excess of exogenous nucleotides is likely to disturb the endogenous occupancy of the Rpt ATPase active sites. Since the conformation of the Rpt subunits is dependent on their nucleotide occupancy and hydrolysis, as the driving force for protein unfolding and substrate translocation, an excess of exogenous ATP is also likely to disturb the endogenous 26S proteasome conformational states. We therefore explored the preparation of human 26S proteasomes in the absence of exogenous nucleotides, revealing unexpected effects on the 26S proteasome peptidase activities and structural integrity driven by exposure to Ca^2+^ or Mg^2^.

We observe that the human 26S proteasome dissociates into its 20S-PC and 19S-RPs sub-complexes when ATP is removed from standard purification buffers containing MgCl_2_, in agreement with the consensus in the field that an excess of ATP (or an analogue) is required for the 26S proteasome stability *in vitro*, as demonstrated in early studies (Eytan et al., 1989; Ganoth et al., 1988; Hough et al., 1987). However, we found that stable 26S proteasomes are obtained in the absence of exogenous nucleotides if Mg^2+^ is also excluded and replaced by EDTA, particularly during the cell lysis. We further investigated the consequences of exposing the 26S proteasome to Mg^2+^, and extended these studies to test if similar effects would be observed upon exposure to the closely related and physiologically highly relevant Ca^2+^. Peptidase activity assays showed that when Ca^2+^ or Mg^2+^ are added to 26S proteasomes prepared by avoiding exposure to these cations, they affect the proteasome activity profile in a similar way, with a distinct modulation of each of the proteasome peptidase activities (Figure 1C). Additionally, we observed that exposure to these cations also disrupts the structural integrity of the 26S proteasomes, leading to the dissociation of its 20S-PC and 19S-RP subcomplexes.

Ca^2+^ is a key intracellular signaling molecule, controlling a wide range of processes including transcription, cell division, apoptosis, synaptic transmission and muscle contraction (Berridge et al., 2000; Clapham, 2007; Petersen et al., 2005). The resting concentration of Ca^2+^ in the cytoplasm and nucleoplasm, where proteasomes are active, is critically maintained at around 100 nM. While the measured amounts of intracellular Mg^2+^ are significant higher, this is found mostly coordinated with other molecules, specially ATP, rather than as free intracellular Mg^2+^. Consequently, under normal physiological conditions 26S proteasomes are exposed to very low concentrations of either free Ca^2+^ or Mg^2+^. However, during purification in the absence of EDTA, 26S proteasomes are likely to be exposed to significantly higher concentrations of Ca^2+^, released from intracellular stores during cell lysis, resulting in its structural instability. To unify the data presented here with previous studies, it is plausible to suggest that exposure to Ca^2+^ or Mg^2+^ may cause either the loss of a still unknown cofactor or a proteasome structural alteration that is somehow recovered *in vitro* by the addition of an excess of exogenous ATP. Overall, the identification of a level of 26S proteasome regulation by Ca^2+^ has significant physiological implications, including in disorders were both proteasomal disfunction and Ca^2+^ signaling dysregulation are observed, such as in neurodegenerative conditions.

### Allosteric communication within the 26S proteasome

While the presence of exogenous ATP (or an analogue) can somehow overcome the 26S proteasome structural instability resulting from exposure to Ca^2+^ or Mg^2+^, they may not fully restore the allosteric communication between 20S-PC and 19S-RPs. Previous studies showed that the chymotrypsin-like activity of conventionally purified 26S proteasomes is enhanced upon binding of ubiquitinated substrates (Bech-Otschir et al., 2009; Nathan et al., 2013). This has been attributed to conformational rearrangements triggered by substrate binding that propagate allosterically onto the 20S-PC proteolytic active sites. However, when model ubiquitinated substrates were added to our 26S proteasomes, prepared without exposure to Mg^2+^ or Ca^2+^, we observed no alteration of their peptidase activities. Nevertheless, the interaction of these proteasomes with the different ubiquitin degron models tested could still be monitored by peptidase activity enhancement, but this required the addition of MgCl_2_ to the reaction buffer used in the activity assays (Figure 2A and C). This suggests that, without exposure to Mg^2+^ or Ca^2+^, the 26S proteasomes retain an active conformation where its peptidase active sites are allosterically uncoupled from substrate binding. Accordingly, cryo-EM analysis revealed that the binding of a ubiquitin degron mimetic had no overall effects on the conformation of the 26S proteasome (Figures 3 and 6). On the other hand, the current data suggests that when the 26S is exposed to Mg^2+^ or Ca^2+^ there is a disruption of its allosteric properties, even in the presence of ATP (or an analogue), which can still be restored, at least to some extent, by structural stabilization induced by substrate binding that can be monitored by peptidase activity enhancement (Figure 7). Importantly, our data indicates that 26S proteasome conformational rearrangements associated with ubiquitinated substrate processing occur downstream of their transition from apo to substrate pre-engagement state, at the ATP dependent steps of unfolding and translocation.

**Figure 7.**
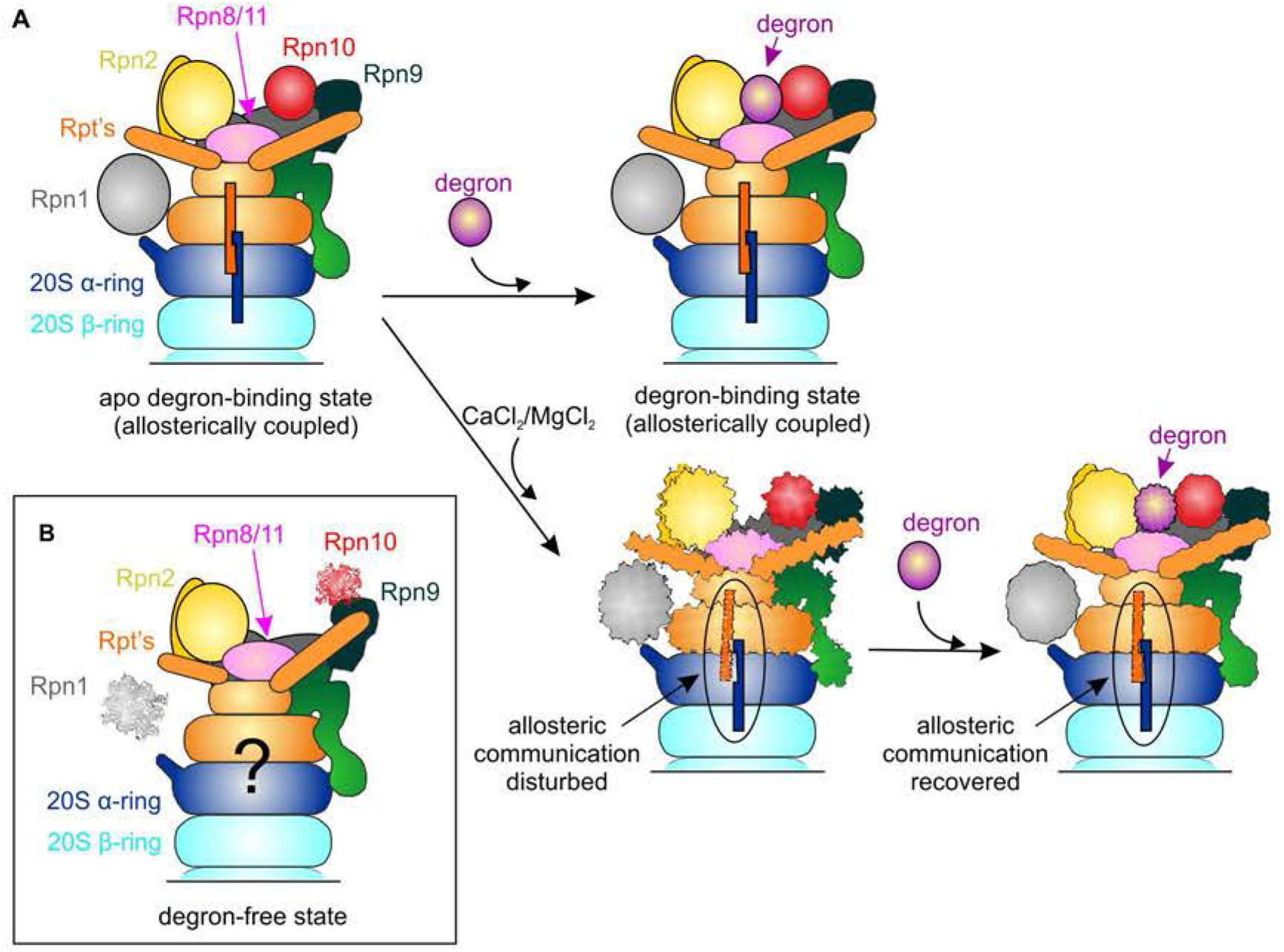
Model of the effects of Ca^2+^ and Mg^2+^ on the 26S proteasome allosteric behavior. **(A)** Representation of the effects of Ca^2+^ and Mg^2+^ on the allosteric communication between the 19S-RP ubiquitin degron binding site and the 20S-PC peptidase activities, shown for the 26S proteasome degron-binding state. The endogenous 26S proteasome allosterically coupled conformation is not altered by degron binding. However, this coupling can be disturbed by exposure to Ca^2+^/Mg^2+^, which is associated with a reduced structural stability of the 26S proteasome. Under these conditions, ubiquitin degron binding can result in a structural stabilization that restores the allosteric communication between the 19S-RP and 20S-PC. **(B)** The allosteric communication between the 19S-RP and the 20S-PC in the 26S proteasome degron-free state cannot be assessed with the current data.

### The 26S proteasome conformations

The cryo-EM analysis of 26S proteasomes in the absence of exogenous nucleotides clearly resolved two conformational states (Figures 3 and 6). Compared with these, any other proteasome conformations that may co-exist in these samples will be highly underrepresented. The cryo-EM analysis of the 26S proteasome in the presence of an optimized ubiquitin degron mimetic, UBg_8UBgg, reveals that only one of the conformations has the degron bound (Figure 3A). In both 26S proteasome states the Rpt hetero-hexamer departs from a closed ring by a cleft associated with the outward displacement of two subunits, Rpt1-Rpt2 and Rpt5-Rpt1 for the degron-free and degron-bound conformations, respectively (Figure 4). In both states these subunits are found in the apo state, with no nucleotides bound, while the remaining four Rpts are nucleotide bound (Figures 4C and S6). Since UBg_8UBgg was added to the 26S proteasome in the absence of exogenous nucleotides, it is extremely unlikely that its binding could result in an endogenous nucleotide transfer between subunits. Furthermore, a control cryo-EM analysis of 26S proteasomes in the absence of degron resulted in the same two conformations being resolved, at a ratio comparable to that observed for the proteasome-UBg_8UBgg sample (Figure 6). These results indicate that the two 26S proteasome states are endogenous and co-exist, with only one able to bind the ubiquitin degron in what appears to be a conformational selection mechanism.

The two proteasome conformations resolved diverge from the S_A_ state found to dominate 26S proteasome samples analyzed by others, in the presence of MgCl_2_ and exogenous nucleotides (Table S2). However, a close comparison of our 26S proteasome structures with those previously reported revealed that the degron-free and degron-binding conformations are related, but not identical, to the E_C2_ and E_D1_ states previously described (Dong et al., 2019), respectively (Figure S7 and Table S2). The main differences between these conformations relate to the nucleotide occupancy and the insertion of the Rpt C-terminal tails into their binding pockets at the 20S-PC surface.

In both the 26S proteasome degron-free conformation and E_C2_ the subunits Rpt6-Rpt3-Rpt4-Rpt5 have the same nucleotide occupancy, with Rpt1 and Rpt2 in the apo state. However, while in the degron-free conformation all Rpt subunits, except Rpt4, have their C-terminal tails inserted into their 20S-PC pockets, in the E_C2_ state the tails for both Rpt1 and Rpt4 are not recovered. Nevertheless, in both states the 20S-PC gate is found in a closed state. This appears to indicate that 20S-PC gate opening is strongly associated with the overall conformation of the Rpts, rather than just the insertion of the Rpt C-terminal tails into their 20S-PC binding pockets. On the other hand, while both the 26S proteasome degron-binding conformation and the E_D1_ state are found to have all their Rpt C-terminal tails inserted into the 20S-PC, except for Rpt4, in the degron-binding conformation Rpt5-Rpt1 are in apo state, while in the E_D1_ state only Rpt5 is found without a nucleotide bound.

It is important to note that, while the S_A_ state, where all the Rpt subunits have ATP bound, dominate in 26S proteasome samples analyzed in the presence of MgCl_2_ and exogenous nucleotides, the E_C2_ and E_D1_ states were resolved in a sample with excess of exogenous nucleotide, but where MgCl_2_ was excluded from the last step of purification and grid preparation, although it was added at earlier stages (Dong et al., 2019). Because the proteasome sample from which E_C2_ and E_D1_ were resolved also had a proteasome substrate added, they were previously interpreted as different steps of substrate processing. Conversely, we show that similar conformations occur with no substrate present.

The data presented here suggests that some of the 26S proteasome conformations previously identified may be related with the environment in which the samples were analyzed, namely exposure to Ca^2+^ (during cell lysis) and/or Mg^2+^, and the presence of saturating concentrations of exogenous nucleotides. In our degron-binding conformation Rtpr5-Rpt1 are found in apo state, while Rpt2-Rpt6-Rpt3-Rpt4 are found bound to ATP. In the degron-free conformation, Rpt1-Rpt2 are found in the apo state, Rpt6-Rpt3 have ATP bound and Rpt4-Rpt5 bind ADP. Notably, this nucleotide occupancy of the 26S proteasome in the degron-free and degron-binding conformations is consistent with earlier biochemical studies indicating that the Rpt subunits function in pairs. According to these studies, in AAA-ATPases bound to the 20S-PC only four subunits can bind ATP at any time, with 3 pairs of adjacent Rpt subunits binding ATP with distinct affinities (Kim et al., 2015; Smith et al., 2011).

Surprisingly, unassigned densities are found at the central pore of the Rpt AAA-ATPase ring in the cryo-EM maps of both 26S proteasome degron-free and degron-binding conformations (Figure S6C). The appearance of these densities is consistent with a polypeptide chain and their strength indicates high occupancy. In each of our structures this polypeptide is surrounded by Rpt subunits, which are Rpt2-Rpt6-Rpt3-Rpt4 in the degron-binding conformation and Rpt6-Rpt3-Rpt4-Rpt5 in the degron-free state (Figure S6C). In both cases, the two Rpt subunits displaced from a closed Rpt ring conformation, which are Rpt1 and Rpt5 in the degron-binding and Rpt1 and Rpt2 in the degron-free states (Figure 4), do not appear to interact with the polypeptide chain. It is important to note that similar densities were also found in the related E_C2_ and E_D1_ conformations (Dong et al., 2019), where they were assigned to an added substrate being processed by the proteasome, but not in the 26S proteasome S_A_ state that dominates in the presence of Mg^2+^ and an excess of nucleotides (Table S2). It can be speculated that these densities in the maps presented here may correspond to endogenous proteasome substrates that are carried over during purification, although an unambiguous density assignment is still required.

### The ubiquitin degron mimetic and its binding to the 26S proteasome

In the cryo-EM map of the degron-binding 26S proteasome the densities for UBg_8UBgg bridge Rpn10 and Rpn2 (Figure 3A), in agreement with previous structural studies (de la Peña et al., 2018; Dong et al., 2019). Rpn10 is the canonical 26S proteasome ubiquitin receptor (Young et al., 1998). The cryo-EM map of the proteasome in its degron-binding conformation has strong densities for Rpn10, as is the case of the equivalent conformation in the control analysis in the absence of degron (Figures 3A and 6A). These are significantly weaker in the maps of the degron-free conformation, suggestive of higher Rpn10 structural flexibility and/or lower occupancy hindering the binding of UBg_8UBgg. The 26S proteasome in the degron-binding state can be assumed to be proteolytically active. On the other hand, while the proteasome degron-free state may correspond to either an inhibited or latent state of the proteasome proteolytic cycle, it cannot be excluded that this may also be proteolytically active against proteins targeted for proteasomal degradation by other non-canonical signals, which are known for their diversity (Ciechanover and Stanhill, 2014).

It has been shown that only two ubiquitin moieties of K48-linked tetra-ubiquitin chains interact with the proteasome, namely ubiquitin-1, proximal to the substrate, and ubiquitin-3 (Thrower et al., 2000). UBg_8UBgg was optimized to mimic the endogenous K48-linked tetra-ubiquitin degron and both have similar proteasome binding affinities (Figure 2D and E), suggesting that the two ubiquitin moieties in UBg_8UBgg are optimally spaced for proteasome interaction. Information on the orientation of the degron in its proteasome binding pocket can be inferred from the biochemical data presented here. N-terminal extensions on di-ubiquitin model degrons had no effect on their interaction with the proteasome, while extending the C-terminus results in a considerable loss of affinity (constructs 21UBg_12UBgg and UBg_12UBg_21, respectively, compared with UBg_12UBg; Figure S2 and Table S1). The endogenous ubiquitin degron is conjugated to substrate proteins by the C-terminus of its proximal ubiquitin, which should face the proteasome degron binding pocket to be positioned for cleavage from the substrate by Rpn11. It is therefore plausible that the N- and C-terminal ubiquitin moieties of UBg_8UBgg mimic ubiquitin-3 and −1 of the endogenous tetra-ubiquitin degron, consistent with C-terminal extensions on degron mimetics disrupting the interaction with the proteasome, albeit with a geometry significantly distinct from that of its conjugation to a substrate lysine residue, resulting in a significantly reduced binding affinity to the 26S proteasome. The data presented here allows the assignment of the degron-binding 26S proteasome conformation as its substrate pre-engagement state, with any proteasome conformational re-arrangements occurring downstream during the substrate processing path, which can now be unambiguously characterized.

### Final remarks

Our results represent a significant contribution towards the full characterization of the human 26S proteasome. The unexpected finding that exposure to Ca^2+^ and Mg^2+^ affects the 26S proteasome activity and destabilizes its structural integrity has substantial consequences for future functional and structural studies on the complex. Notably, the proteasome regulation by Ca^2+^, a key intracellular regulatory signal, may have significant physiological consequences, and it may underlie proteasome disfunction in physiological conditions associated with Ca^2+^ signaling dysregulation, including in neurodegenerative conditions. The data presented here is likely to ultimately pave the way to important and improved translational applications, as the proteasome is already a recognized therapeutic target for diverse serious medical conditions.

Our demonstration that stable 26S proteasomes can be prepared in the absence of exogenous nucleotides, when not exposed to Ca^2+^ or Mg^2+^, together with the characterization of the two proteasome conformations resolved and their interaction with a ubiquitin degron, opens substantial new opportunities in fundamental studies on the still significantly elusive 26S proteasome function and regulation.

## Acknowledgements

We are grateful to E. Morris for his contribution to the preparation of cryo-EM grids and for the critical reading of the manuscript. We thank J. Grimmett and T. Darling for computing support. This study was also supported by the Biophysics and Electron Microscopy Facilities at the MRC-LMB. The work was funded by the Medical Research Council grant MC_UP_1201/5 to P.dF.

## Author contributions

M.K. and P.dF. designed the experiments; M.K., J.T. and A.T.R. optimized the 26S proteasome purification protocol; M.K. prepared and characterized all degrons tested; M.K., P.A. and P.dF. did the cryo-EM analysis, M.K., P.A., A.T.R and P.dF. did the model building. M.K. and P.dF. wrote the manuscript and all authors critically reviewed it and agreed with its submitted version.

## Competing interests

The authors declare no competing interests

## Materials & Correspondence

Requests should be addressed to P.dF..

## Data Availability

EM maps have been deposited in the Electron Microscopy Data Bank under accession codes XXX. The atomic models have been deposited in the Protein Data Bank under accession numbers XXX

## METHODS

### Purification of human 26S proteasomes

Human embryonic kidney cells adapted to suspension culture (HEK293F) were grown in Free Style 293 Expression Medium (Thermo Fisher Scientific) and harvested at a density of 3 million cells/ml. Cells were collected by centrifugation (10 min, 4000 x g) and washed twice (10 min, 1500 x g) in standard buffer (50 mM Tris-HCl pH 7.4, 50 mM NaCl, 5 mM MgCl_2_, 10% glycerol, 5 mM ATP, 0.25 mM TCEP) for conventional 26S proteasome purification, or exogenous nucleotide depleted buffer (50 mM HEPES pH 7.5, 50 mM NaCl, 5mM EDTA, 10% glycerol, 0.25 mM TCEP) for the purification of 26S proteasomes in the absence of added nucleotides and avoiding exposure to Ca^2+^/Mg^2+^. Resulting washed pellets were frozen and stored at −80°C.

In a typical purification, a 25 g pellet of HEK293F cells was resuspended in 75 ml of either standard or exogenous nucleotide depleted buffer. Cells were disrupted by using a dounce homogeniser and the cell lysate was briefly centrifuged (10 min, 12000 x g). After centrifugation, the supernatant was filtered, mixed with human strep-tagged Rad23b-UBL (adapted from a previously described protocol (Besche et al., 2009)) and incubated for 1h at 4°C. 0.8 mg of avidin was added just prior to affinity chromatography. A 5 ml Strep-tactin Superflow Plus column (Qiagen) was equilibrated with 50 mM Tris-HCl pH 7.4, 50 mM NaCl, 5 mM MgCl_2_, 10% glycerol, 5 mM ATP, 0.25 mM TCEP, or 50 mM HEPES pH 7.5, 50 mM NaCl, 5 mM EDTA, 10% glycerol, 0.25 mM TCEP, and bound proteins were eluted with 3.5 mM d-desthiobiotin (Sigma) in the equilibration buffer (Besche and Goldberg, 2012; Besche et al., 2009; Peth et al., 2013). The 26S proteasome containing fractions, eluted from the affinity chromatography column, were separated on a glycerol gradient (20 h, 124000 x g). Double-capped 26S proteasomes were concentrated in 100 kDa Amicon Ultra (Merck) centrifugal filter (45 min, 1500 x g).

For high resolution cryo-EM, 100 μl of the concentrated 26S proteasome sample (alone or mixed with 90 μM of model degron) was further purified by size exclusion chromatography using a Superose 6 PC 3.2/30 column (GE Healthcare), equilibrated with 50 mM HEPES pH 7.5, 50 mM NaCl, 5 mM EDTA, 2% glycerol, 0.25 mM TCEP.

### Proteasome peptidase activity assays

The three 26S proteasome peptidase activities were measured using specific 7-amino-4-methylcoumarin (AMC) tagged fluorogenic peptides, namely Suc-L-L-V-Y-AMC (chymotrypsin-like activity), Boc-L-R-R-AMC (trypsin-like activity) and Z-L-L-E-AMC (caspase-like activity) (Boston Biochem). The proteasome samples were incubated with 50 μM AMC-peptide in the respective purification buffers (standard or the exogenous nucleotide depleted buffer) for 30 min at 25°C.

For the measurements of the Ca^2+^/Mg^2+^ effects, the 26S proteasome samples were incubated with 50 μM AMC-peptide in 50 mM HEPES pH 7.5, with different concentrations of MgCl_2_ or CaCl_2_ added. Fluorescence was measured using a PHERAstar FS plate reader (BMG Labtech) with an excitation wavelength of 350 nm and an emission wavelength of 450 nm. The gain was adjusted to a 1 μg/ml chymotrypsin containing sample. All measurements were repeated at least 3 times.

### 26S proteasome degradation of a folded model substrate

A previously described model folded substrate (Bhattacharyya et al., 2016) was composed of an N-terminal Rad23b-UBL domain, which consisted of residues 1-79 of human Rad23b, followed by a GSGGSGSG linker, a SNAP-tag domain, a 40 amino acid unstructured region from human cytochrome b_2,_ and a C-terminal hexahistidine tag. This sequence was cloned in a pET11 vector, expressed in *E. coli* BL21 Gold expression strain and purified as previously described (Bhattacharyya et al., 2016). In brief, cells were grown at 37°C to an OD_600_ of 0.6, then induced with 0.5 mM IPTG for a period of 3 h at 28°C. Cells were harvested and resuspended in 50 mM sodium phosphate pH 8.0, 300 mM NaCl, 10 mM imidazole, and 1 mM DTT and a protease inhibitor cocktail (Roche). Cells were lysed by sonication and incubated for 10 min at 25°C with DNase I (Sigma) and 4 mM MgCl_2_.

The lysate was centrifuged (20 min, 48000 x g), the supernatant was loaded on a HisTrap HP 5 ml column (GE Healthcare) and the His-tagged protein was eluted with four column volumes of 50 mM sodium phosphate pH 8.0, 300 mM NaCl, 250 mM imidazole, and 1 mM DTT. Eluted fractions were analyzed by SDS-PAGE and the purest fraction was used for fluorescent dye-labelling. 5 mM of purified protein was combined with 2 mM DTT and 10 mM SNAP-Surface Alexa Fluor 546 (New England Biolabs) in PBS for 60 min at 30°C. Free dye was removed by size exclusion chromatography using a HiPrep 16/60 Sephacryl S 100 HR column (GE Healthcare) equilibrated with PBS buffer. Fractions containing the dye-labelled protein were detected by in-gel fluorescence imaging with an Amersham Typhoon (GE Healthcare) laser scanner.

For the fluorescence anisotropy measurements, the 26S proteasome at a concentration of 50 nM was mixed with 20 nM Alexa Fluor 546-labelled folded model substrate (Bhattacharyya et al., 2016). Reactions were carried out in 50 mM Tris-HCl pH 7.4, 5 mM MgCl_2_, 2 mM ATP 2% glycerol, 1mM DTT, and 0.2 mg/mL BSA at 37°C. Measurements were performed using a PHERAstar FS plate reader (excitation at 535 nm and emission at 580 nm). A polarization reading was taken every 30 min. Anisotropy values (r) were calculated from the polarization readings (P) using the equation r = (2P)/(3 – P).

### Model degrons expression and purification

The model degron constructs were generated from the synthetic gene of the longest construct (ubiquitin - 52 amino acid loop - ubiquitin (UB52UB)) (Figure S2). The N-terminal ubiquitin in all the constructs had either a G76 deletion, or the G75 and G76 residues mutated to V75 and V76. Constructs were named accordingly, by adding the mutation letter to the degron name (for example UBg_20UBgg and UBvv26UBgg respectively). Constructs were built in pET11 vectors and expressed in *E. coli*. Modifications were introduced using *in vivo* assembly (IVA) cloning (García-Nafría et al., 2016). All model degrons were expressed soluble and purified using a protocol similar to that used for ubiquitin purification (Pickart and Raasi, 2005). In brief, cells were grown at 37°C to an OD_600_ of 1.0 and induced with 0.4 mM IPTG for 3 h. The cells were then collected and resuspended in lysis buffer (50 mM Tris-HCl pH 7.4) containing a protease inhibitor cocktail (Roche), lysed by sonication and incubated for 10 min at 25°C with DNase I (Sigma) and 4 mM MgCl_2_. The lysate was centrifuged (20 min, 48000 x g) and soluble proteins in the supernatant were precipitated by adjusting the pH to 4.5. The protein solution was centrifuged (10 min, 48000 x g) and the supernatant containing the model degron was dialyzed overnight, at 4°C, to 50 mM ammonium acetate pH 4.5. The solution was filtered and the model degrons were loaded on a HiTrap SP HP 5 ml column (GE Healthcare) in 50 mM ammonium acetate pH 4.5. Proteins were eluted using a linear gradient (from 0% to 100% in 20 column volumes) of elution buffer (50 mM ammonium acetate, 1 M NaCl). Peak containing fractions were pooled, concentrated and further purified on a HiPrep 16/60 Sephacryl S 100 HR column (GE Healthcare) in 50 mM Tris-HCl pH 7.4. The purified proteins were concentrated and flash frozen in small aliquots for storage at −80°C.

### Assessment of 26S proteasome interaction with model degrons

An initial assessment of the interaction of model degrons with the 26S proteasome was performed by monitoring the proteasome chymotrypsin-like activity in the presence of 10 μM of each degron, measured as for the proteasome peptidase activity assays described above except for the assay buffer that was replaced by 50 mM Tris-HCl pH 7.4, 20 mM MgCl_2_, 1 mM DTT and 0.2 mg/mL BSA. Control experiments were performed with commercial human 20S-PC (Enzo Biochem) and endogenous ubiquitin chains (Enzo Biochem).

Similar assays were used to quantitatively determine the binding affinity of the degrons to the 26S proteasome. Here, a multi-substrate reaction rate equation was used to describe the binding of activator (degron) to the proteasome in the presence of substrate (peptide), where the concentration of the substrate is kept constant and that of the activator varied. A set of activity assays was performed at different fixed concentrations of each of the model degrons tested (serial dilutions from 10 μM to 10 nM). Fluorescence was monitored as described above.

### Cryo-electron microscopy grid preparation and data collection

The 26S proteasome-UBg_8UBgg complex was used for grid preparation directly after size exclusion chromatography co-purification, with additional 1 μM (5 fold higher that the measured binding affinity) of UBg_8UBgg to ensure the saturation of the 26S proteasome. This sample was applied on 400 gold mesh Quantifoil R1.2/1.3 electron microscope grids (Quantifoil Micro Tools), freshly coated in house with an additional thin layer of carbon and glow discharged. The grids were plunged-frozen into liquid ethane using a Vitrobot Mark IV (Thermo Fisher Scientific), operated at 22°C and 95% humidity. The grids were transferred into a Thermo Fisher Scientific Titan Krios electron microscope (Thermo Fisher Scientific), at the MRC-LMB facility, operating at 300 keV and a nominal magnification of x75,000, resulting in a calibrated sampling of 1.04 Å per pixel at the image level. Images were recorded with EPU software using a Falcon III detector operating in counting mode, at an electron dose per pixel of ∼0.5 e^-^/s, with 60 s exposures saved as 75 frame movies with an evenly distributed electron dose (Figure S4B). The recorded images had a measured defocus range of −0.6 to −3.8 μm.

### Single particle analysis

The overall image processing workflow is shown in Figure S5 (see also Table S3). A total of 5494 recorded movies of the 26S proteasome, with excess of UBg_8UBgg, were recorded. For each movie, frames 3 to 75 were aligned, dose-weighted and summed using MotionCorr whole-image motion correction software (Li et al., 2013). The summed images were used for contrast transfer function estimation in Gctf (Zhang, 2016). 4505 images were selected for further processing, based on ice quality, image contrast and the recovery of isotropic Thon rings in their power spectra. Particle auto-picking was performed with Gautomatch (http://www.mrc-lmb.cam.ac.uk/kzhang/Gautomatch), resulting in a dataset of 822,102 particles that were extracted into 640×640 pixel boxes. The particles were down-sampled (160×160 pixels) and subjected to multiple rounds of two-dimensional (2D) classification in RELION 3.0 (Scheres, 2012). The initial cycles of 2D classification were performed with a large search range (100 Å) to optimize the centering of the particles. The particles grouped in 2D class-averages corresponding to uncapped complexes, remaining false-positive picked particles and poorly centered proteasome particles were removed after subsequent rounds of 2D classification, yielding a dataset of 132,244 particles. Further analysis was performed with the selected particles at their original sampling. An initial 3D refinement, with applied C2 symmetry, was performed using a double-capped 26S proteasome cryo-EM map, previously obtained in-house and Fourier low-pass filtered to 20 Å. The resulting alignment parameters of each particle were used for symmetry expansion, where each of the two C2 symmetry related proteasome halves was treated as an independent asymmetric particle (equivalent to a dataset of 264,488 asymmetric units). The resulting particle data-set obtained was subjected to 3D classification into 5 classes, with no symmetry imposed and using a soft-edged mask comprising one 19S-RP and half of the 20S-PC. Two of the resulting classes corresponded to well resolved 26S proteasome complexes. The particle components of each of these classes were imported to cisTEM (Grant et al., 2018) for final rounds of 3D classification and refinement, focused on the proteasome 19S-RP. In a final step of the analysis, signal subtraction followed by focused 3D classification of the less recovered regions of the map of the proteasome degron-binding conformation (namely focusing on Rpn1 and the degron/Rpn10 regions of the map) was performed in Relion, but resulted in no significant improvements. The resolution of the final maps was estimated by Fourier shell correlation, as implemented in cisTEM (Figure S4D).

The image processing strategy followed for the 26S proteasome-UBg_8UBgg sample was adapted to analyze a control image data-set recorded of the human 26S proteasome, in the absence of exogenous nucleotides, in its apo state (in the absence of substrates or degron-mimetics). Here 5401 images were recorded, from which 4796 were selected for analysis. Particles were picked using references corresponding to half-proteasome, resulting in an initial data-set of 1,032,174 picked 26S proteasome asymmetric units. After particle picking, removal of false positives and uncapped complexes, a dataset of 352,732 asymmetric units was obtained. These were imported into cisTEM for preliminary 3D refinement followed by 3D classification into 3 classes. One of the classes corresponded to complexes that were either uncapped or were bound to uncomplete 19S-RPs. The other two 3D classes were found to correspond to conformations equivalent to those resolved in the presence of an excess of UBg_8UBgg (Figures 6 and S5), and were not further analyzed.

### Protein model building

The maps of the two 26S proteasome conformations resolved in the presence of an excess of UBg_8UBgg were used for model building, based on previously published human and yeast 26S proteasome cryo-EM structures (PDB IDs: 5t0j, 6fvy, 6msg, 6msh), using real-space refinement in Coot (Emsley et al., 2010) and Phenix (Afonine et al., 2012). The models were manually corrected using the amino acid side chain densities clearly resolved in the maps. When necessary, the maps were sharpened with B-factors to assist model building. The final atomic models were validated using MolProbity (Chen et al., 2010) (Table S3).

### Graphic representations

The representations of the proteasome structures shown in Figures 3, 6 and S4 were created using UCSF Chimera (Pettersen et al., 2004), while the remaining structure representations in the manuscript were created using the Pymol Molecular Graphics System, Schrödinger, LLC. The cryo-EM maps shown in Figure S4C were colored according to local resolution as estimated using ResMap (Kucukelbir et al., 2014). The cryo-EM maps used for the preparation of the figures are unmasked, unsharpened and are Fourier low-pass filtered to 3.6 Å.

## Notes

### Competing Interest Statement

The authors have declared no competing interest.

