## Supplemental Figure 1 for "New insights into the human 26S proteasome function and regulation"

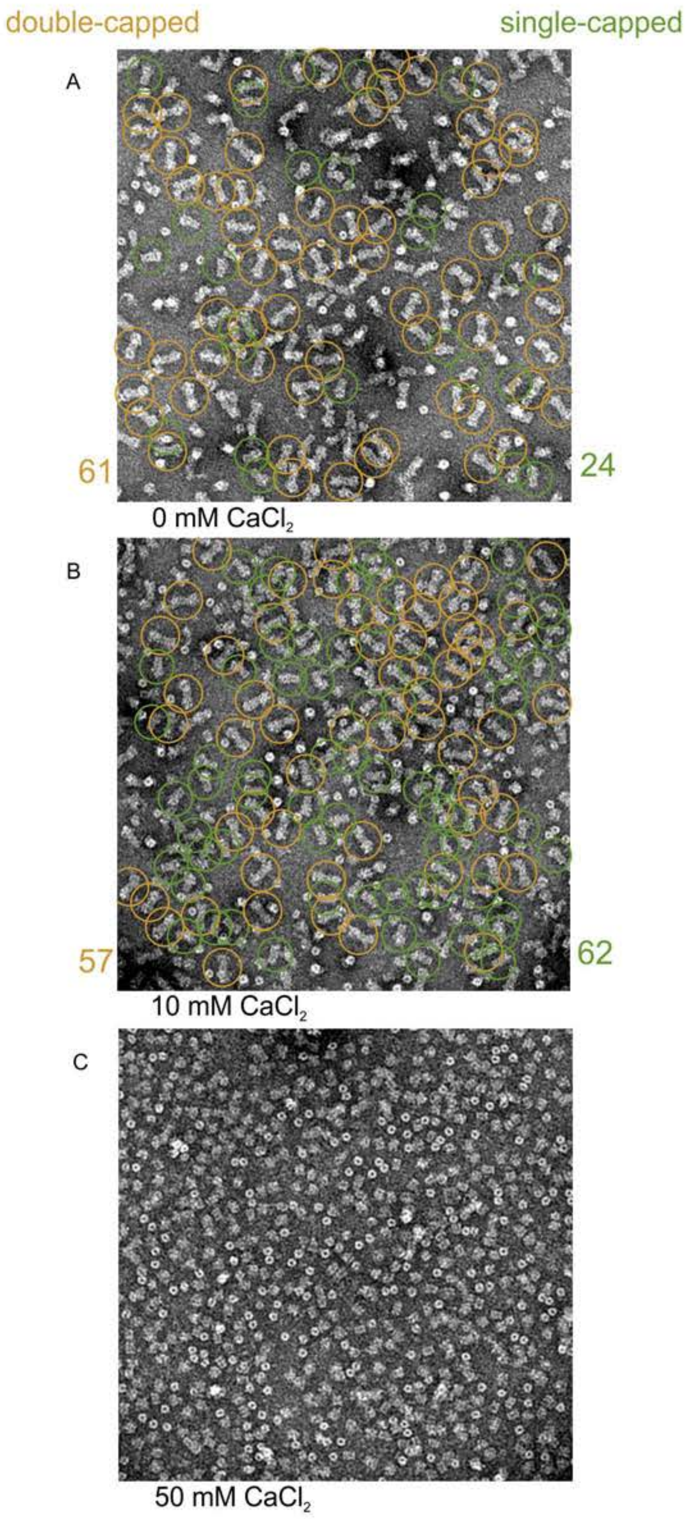


**Figure S1 | The effects of $CaCl_2$ on the stability of the 26S proteasome assembly.** Micrographs of a negatively stained 26S proteasome sample after incubation, for 2 hours at room temperature, with (**A**) 0 mM, (**B**) 10 mM or (**C**) 50 mM of $CaCl_2$. In (**A**) and (**B**) a micrograph is shown with clearly distinguishable double-capped (orange circles) or single-capped (green circles) complexes indicated, with their totals (color coded) shown on the sides of the panels. A ratio of ~5:2 double- to single-capped complexes is observed in the absence of $CaCl_2$, while this changes to closer to ~1:1 in the presence of 10 mM $CaCl_2$. In (**C**) virtually all 26S proteasomes appear dissociated into 20S-PC and 19S-RP.
