## Supplemental Figure 2 for "New insights into the human 26S proteasome function and regulation"

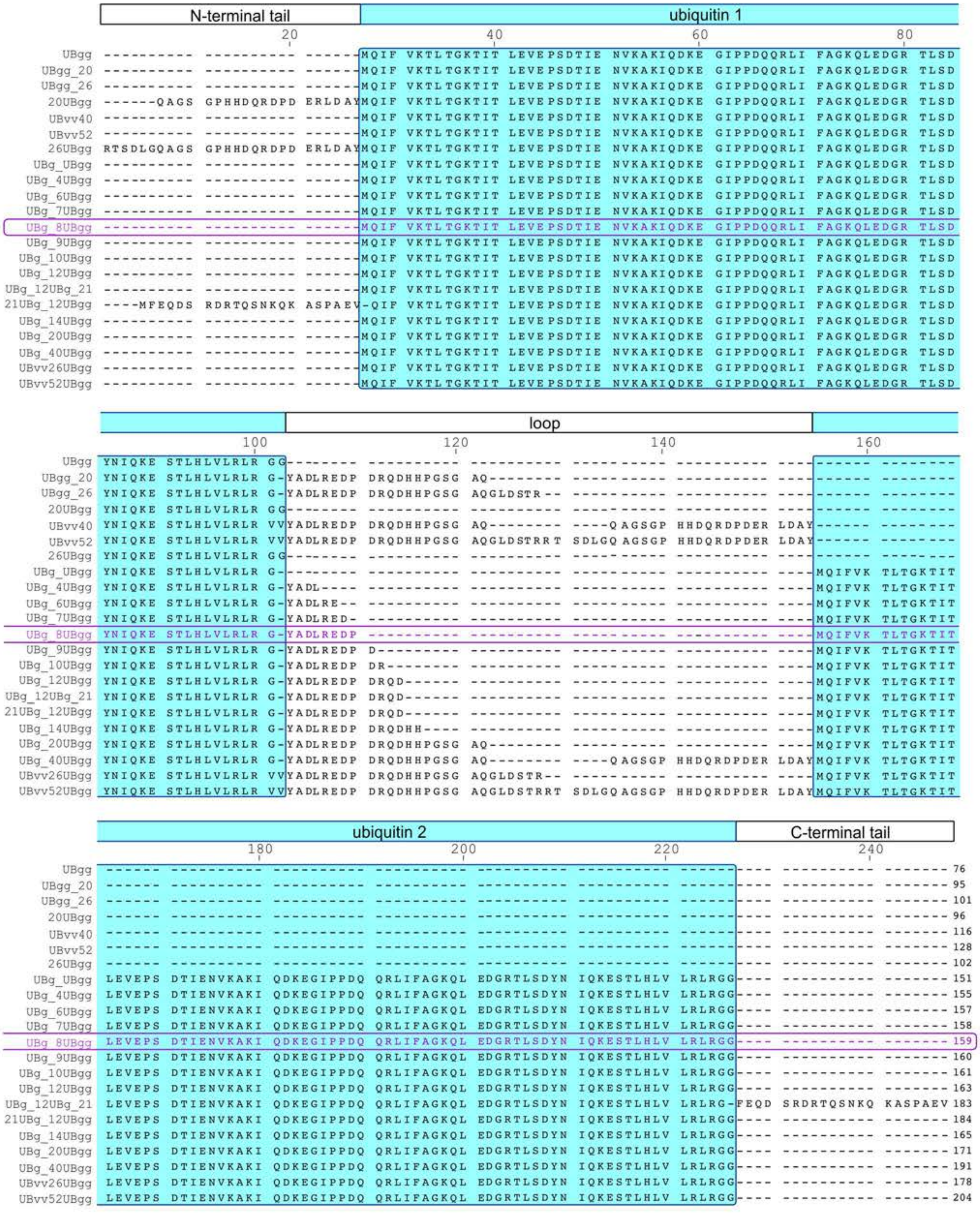


**Figure S2 | Amino acid sequence of ubiquitin derived constructs designed to identify an optimal 26S proteasome ubiquitin-degron mimetic.** The constructs were designed to comprise one or two ubiquitin moieties (highlighted in cyan) flanked by unstructured loops or extensions of different lengths. The sequence of UBg_8UBgg, identified as the best K48 linked tetra-ubiquitin chain mimetic, is shown in purple. The different domains of the constructs are indicated above the sequences.
