## Supplemental Figure 3 for "New insights into the human 26S proteasome function and regulation"

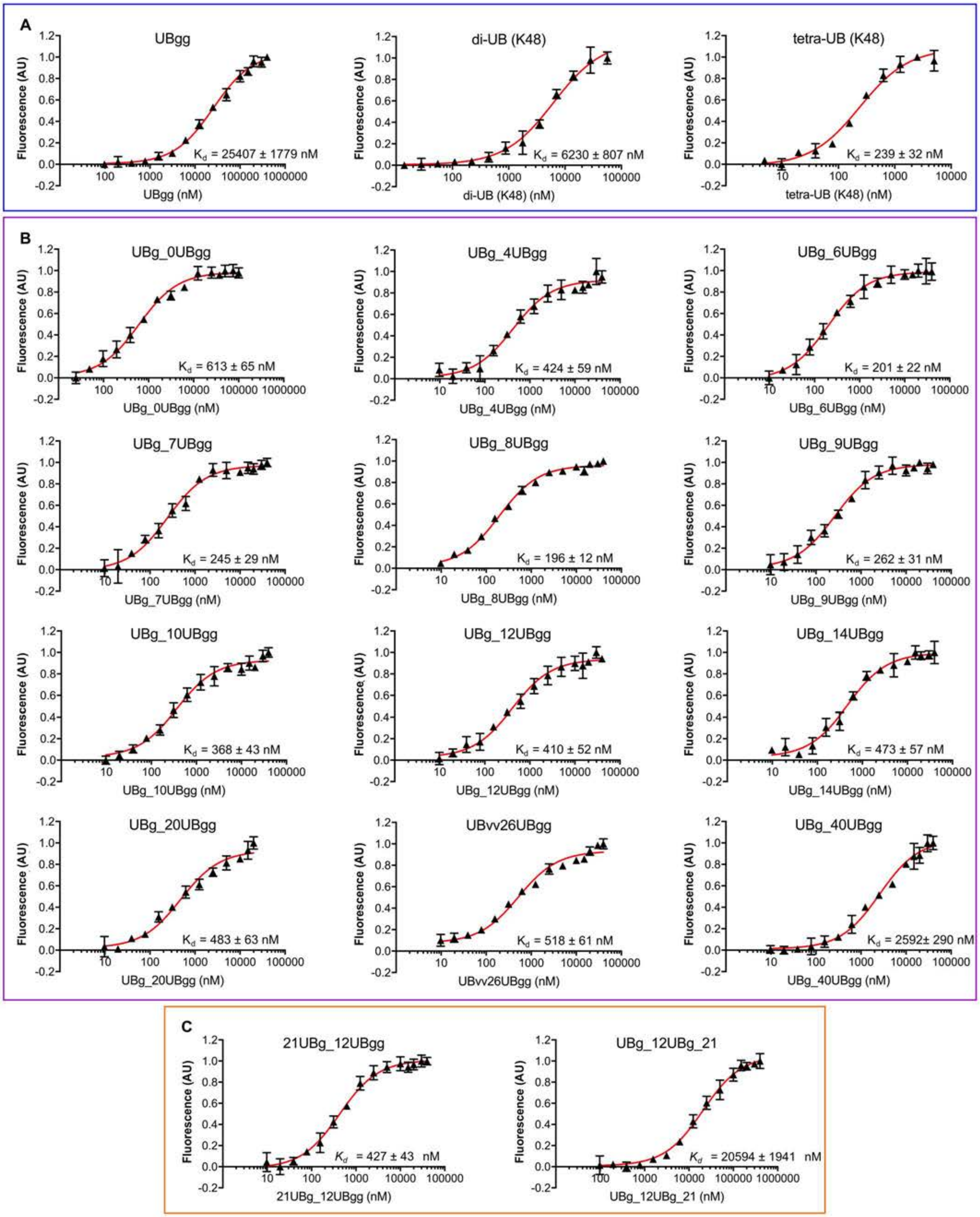


**Figure S3 | Measurement of the binding affinities of model degrons to the human 26S proteasome.** **(A)** Binding affinity of endogenous ubiquitin forms, used as control. **(B)** Binding affinity of constructs formed by two ubiquitin moieties linked by loops of different lengths. **(C)** Measurements used to identify the effects of N- and C-terminal extensions on the model degron binding to the 26S proteasome. Error bars represent the standard deviation from the average of all measurements.
