## Supplemental Figure 4 for "New insights into the human 26S proteasome function and regulation"

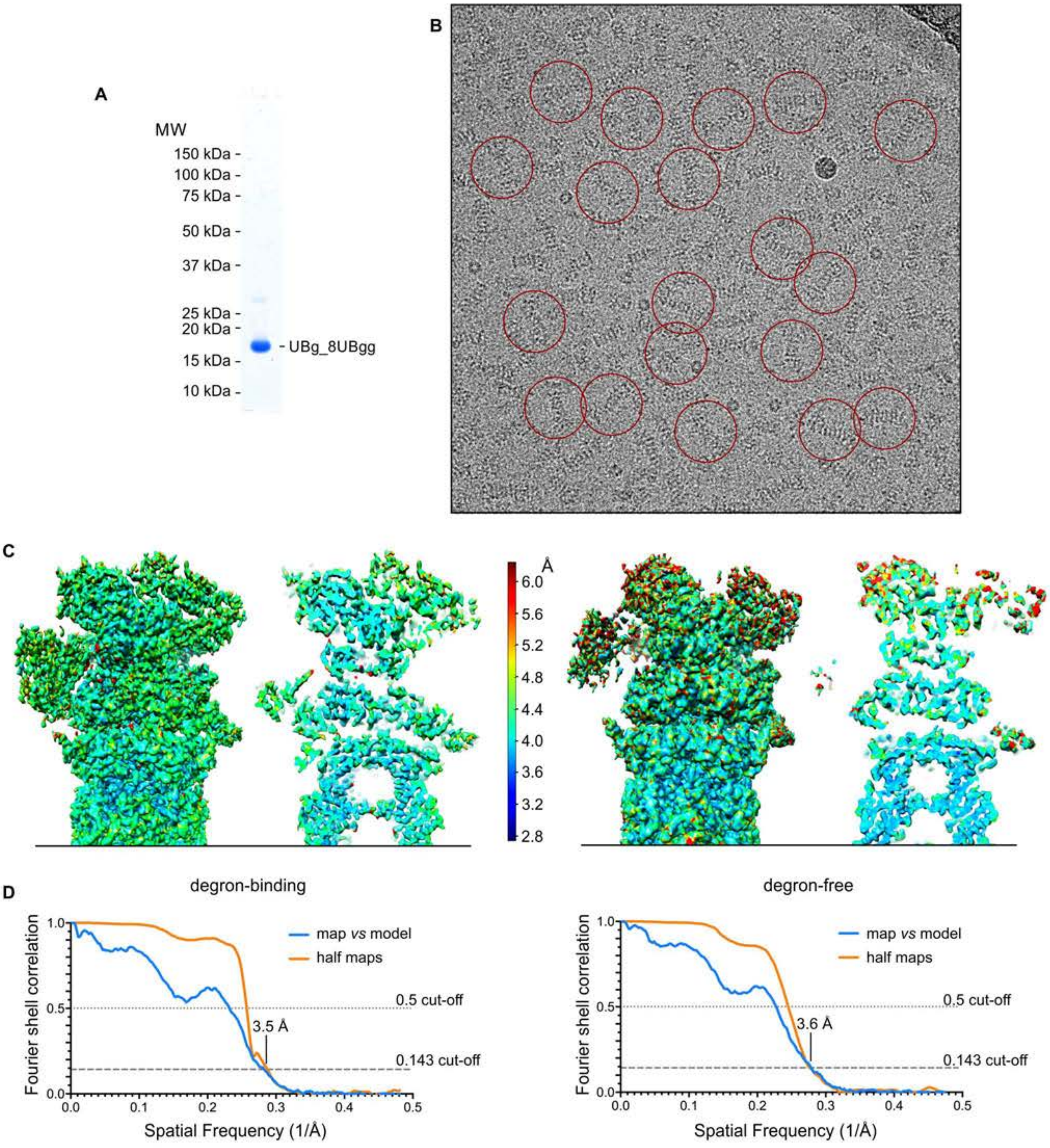


**Figure S4 | Cryo-EM analysis of human 26S proteasomes, not exposed to $Ca^{2+}$ or $Mg^{2+}$ and in the absence of exogenous nucleotides, with added UBg_8UBgg. a,** SDS-PAGE of the purified UBg_8UBgg. **b,** Cryo-EM image of the 26S proteasome sample with UBg_8UBgg. Some of the double-capped 26S proteasome particles in the image are circled. **c,** Surface representation and inner section of the cryo-EM maps of the 26S proteasome degron-binding (left) and degron-free (right) states color coded according to local resolution, as indicated. **d,** Fourier shell correlation curves determined for the final cryo-EM maps of the 26S proteasome degron-binding (left) and degron-free (right) states.
