## Supplemental Figure 5 for "New insights into the human 26S proteasome function and regulation"

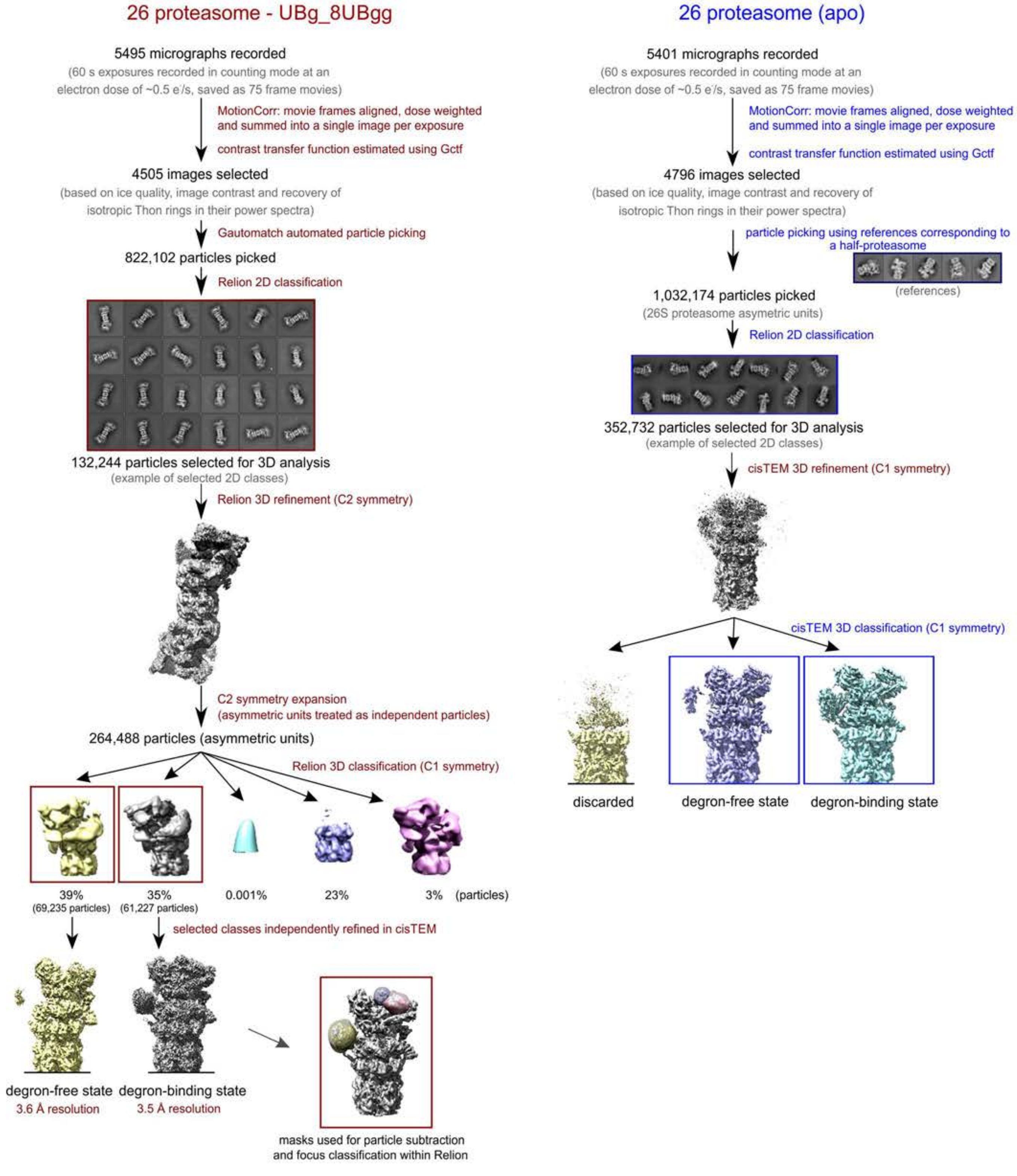


**Figure S5 | Schematic representation of the cryo-EM single-particle analysis workflows.** 26S proteasome samples with added UBg_8UBgg (left) or in the apo state (right) were analysed. The main steps include frame alignment and contrast transfer function determination, 2D classification, initial 3D refinement, 3D classification and the independent refinement of selected classes. Particle subtraction and focused 3D classification was performed for the degron-bound map obtained for the analysis of the 26S proteasome-UBg_8UBgg complex only, using masks focused on Rpn1 (yellow) or combined Rpn10 and degron binding site (pink and purple).
