## Supplemental Figure 6 for "New insights into the human 26S proteasome function and regulation"

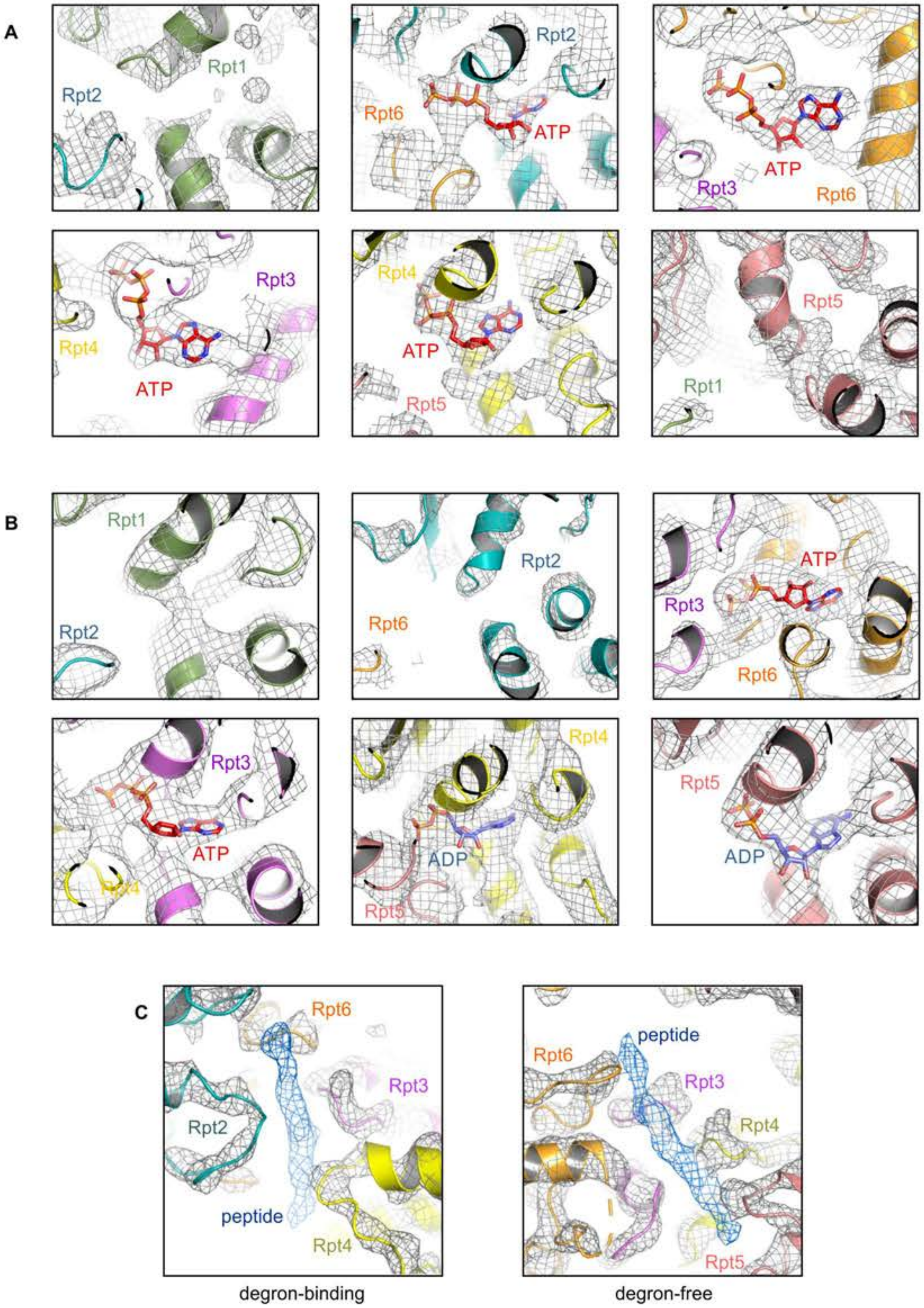


**Figure S6 | Structural details of the 26S proteasome Rtp subunits.** Rpt nucleotide binding sites of the human 26S proteasome degron-binding **(A)** and degron-free **(B)** states. The cryo-EM maps clearly show the presence or absence of bound nucleotides at each binding site (see also Figure 4). The cryo-EM maps are represented as mesh, the protein models as cartoon and the fitted nucleotides as sticks. (**C**) Close-up views of the cryo-EM maps of the degron-binding (left panel) and degron-free (right panel) states of the human 26S proteasome, showing an unassigned peptide density (blue mesh) closely associated with the Rpt subunits (protein model shown as cartoon). In all panels the cryo-EM maps are shown as mesh.
