## Supplemental Figure 7 for "New insights into the human 26S proteasome function and regulation"

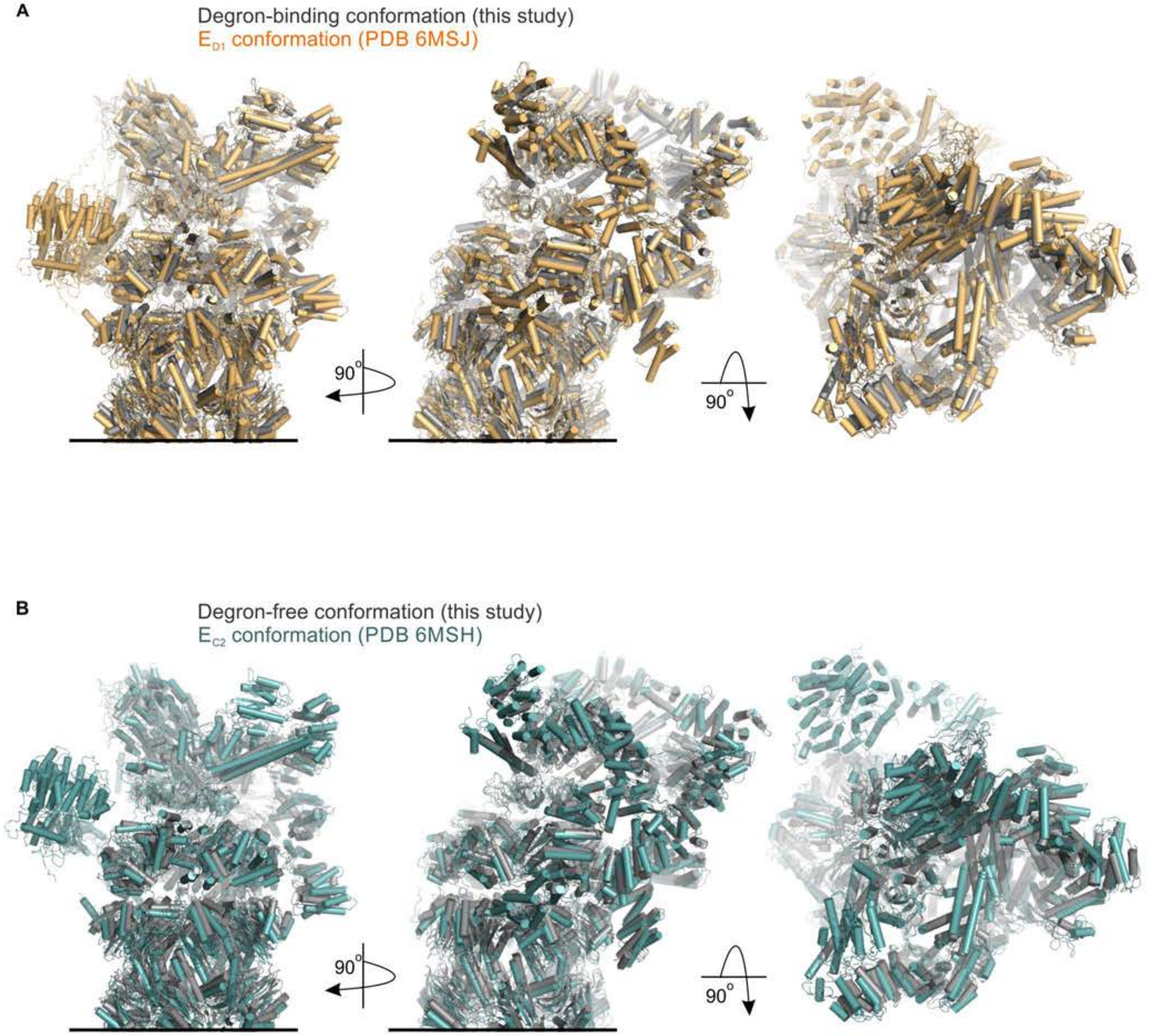


**Figure S7 | Comparison of 26S proteasome conformations.** The degron-binding (**A**) and degron-free (**B**) conformations of the 26S proteasome described here (shown in grey) were compared with those previously described and were found to be closely related, and are shown superimposed, to the $E_{D1}$ (orange) and $E_{C2}$ (cyan) states, respectively (see also, Table S2).
