## Supplemental Table 1 for "New insights into the human 26S proteasome function and regulation"

**Table S1 | Binding affinities of model degrons to the 26S proteasome.**

| Degron | $K_d$ (nM) | R-squared | Loop length (Å) |
|---|---|---|---|
| UBgg | 25407 ± 1779 | 0.9918 | N/A |
| di-UB (K48) | 6230 ± 807 | 0.9822 | N/A |
| tetra-UB (K48) | 239 ± 32 | 0.9821 | N/A |
| UBg_20 | >25000 | N/A | N/A |
| UBg_26 | >25000 | N/A | N/A |
| UBvv40 | >25000 | N/A | N/A |
| UBvv52 | >25000 | N/A | N/A |
| 20UBgg | >25000 | N/A | N/A |
| 26UBgg | 24238 ± 2298 | 0.9876 | N/A |
| UBg_0UBgg | 613 ± 65 | 0.9793 | 0 |
| UBg_4UBgg | 424 ± 59 | 0.963 | 14 |
| UBg_6UBgg | 201 ± 22 | 0.9767 | 22 |
| UBg_7UBgg | 245 ± 29 | 0.9724 | 25 |
| UBg_8UBgg | 196 ± 12 | 0.9925 | 29 |
| UBg_9UBgg | 262 ± 31 | 0.9725 | 32 |
| UBg_10UBgg | 368 ± 43 | 0.974 | 36 |
| UBg_12UBgg | 410 ± 52 | 0.9694 | 43 |
| UBg_14UBgg | 473 ± 57 | 0.9728 | 50 |
| UBg_20UBgg | 483 ± 63 | 0.9706 | 72 |
| UBvv26UBgg | 518 ± 61 | 0.9738 | 94 |
| UBg_40UBgg | 2592 ± 290 | 0.9792 | 144 |
| UBvv52UBgg | 3648 ± 270 | 0.9897 | 187 |
| 21UBg_12UBgg | 427±43 | 0.9806 | 43 |
| UBg_12UBg_21 | 20594±1941 | 0.9848 | 43 |
| UBg_8(polyA)UBgg | 261± 36 | 0.9787 | 29 |

The $K_d$ values were obtained from three independent assays using 50 nM 26S proteasome. The theoretical length of the different loops between the ubiquitin moieties of the degron were calculated assuming that each amino acid has a length of 3.5 Å and that the loops are completely unstructured and extended.
