## Supplemental Table 2 for "New insights into the human 26S proteasome function and regulation"

**Table S2 -** Summary of key features of human 26S proteasome structures, available for the complete complex and resolved at resolutions better than 4Å.

| *Reference* | *Huang* et al., *2016* | *Chen* et al., *2016* | *Zhu* et al., *2018* | *Dong* et al., *2019* | | | | | | *This manuscript* | |
|---|---|---|---|---|---|---|---|---|---|---|---|
| *$MgCl_2$ concentration on grid* | 5 mM | 1 mM | 1 mM | none (5 mM added to buffer used in cell lysis) | | | | | | none | |
| *nucleotide concentration on grid* | 5 mM ATP | 3 mM ATP | 1 mM ATP-γ-S | 1 mM ATP | | | | | | none | |
| *substrate or substrate mimetic added* | no | no | no | yes | | | | | | yes | |
| *proteasome conformation* | $S_A$-like | $S_A$ | $S_A$ | $E_{A1}$ | $E_{A2}$ | $E_B$ | $E_{C1}$ | $E_{C2}$ | $E_{D1}$ | $E_{D2}$ | degron-free ($E_{C2}$-like) | degron-binding ($E_{D1}$-like) |
| *PDB* | 5GJR | 5T0C | 5VFS | 6MSB | 6MSD | 6MSE | 6MSG | 6MSH | 6MSJ | 6MSK | XXX | XXX |
| *EMDB* | 9512 | 8333 | 8666 | 9216 | 9217 | 9218 | 9219 | 9220 | 9221 | 9222 | XXX | XXX |
| *resolution (Å)* | 3.5 | 3.8 | 3.6 | 3.0 | 3.2 | 3.3 | 3.5 | 3.6 | 3.3 | 3.2 | 3.6 | 3.6 |
| *20S-PC gate* | closed | closed | closed | closed | closed | closed | closed | closed | open | Open | closed | open |
| *Rpt's with nucleotide bound* | Rpt1-6 | Rpt1-6 | Rpt1-6 | Rpt1-6 | Rpt1-6 | Rpt1-5 | Rpt1/3-6 | Rpt3-6 | Rpt1-4/6 | Rpt1-3/5/6 | Rpt3-6 | Rpt2-4/6 |
| *Rpt's in apo state* | - | - | - | - | - | Rpt6 | Rpt2 | Rpt1/2 | Rpt5 | Rpt4 | Rpt1/2 | Rpt1/5 |
| *Rpt's displaced from central pore* | Rpt6 | Rpt6 | Rpt6 | Rpt6 | Rpt6 | Rpt6 | Rpt1/2 | Rpt1/2 | Rpt5 | Rpt4 | Rpt1/2 | Rpt1/5 |
| *Rpt's C-tail inserted into 20S-PC pockets* | Rpt3/5 | Rpt3/5 | Rpt3/5 | Rpt3/5 | Rpt3/5 | Rpt2/3/5 | Rpt2/3/5/6 | Rpt2/3/5/6 | Rpt1-3/5/6 | Rpt1-3/5/6 | Rpt1-3/5/6 | Rpt1-3/5/6 |
| *% of particles from the total in final maps* | 100% | 60% | 53% | 8% | 6% | 19% | 9% | 6% | 23% | 28% | 47% | 53% |
