## Supplemental Table 3 for "New insights into the human 26S proteasome function and regulation"

**Table S3 | Cryo-EM data collection parameters and MolProbity model validation for the analysis of the human 26S proteasome in the presence of UBg_8UBgg.**

| **Data collection parameters** | **class 1** | **class 2** |
|---|---|---|
| Given name | degron-bound (DB) | degron-free (DF) |
| Microscope | Titan Krios | |
| Camera | Falcon III (counting mode) | |
| Voltage | 300 | |
| Magnification | 75000 | |
| Total dose $e^-/Å^2$ | 30 | |
| Defocus rage (µm) | -0.6 to -3.8 | |
| Calibrated pixel size (Å) | 1.04 | |
| Micrographs collected | 5495 | |
| Selected micrographs | 4505 | |
| Number of particles (after symmetry expansion) | 264488 | |
| Refined particles | 178433 | |
| Symmetry | C1 | |
| Particles in final classes | 61227 | 69235 |
| Map resolution (Å) | 3.5 | 3.6 |

| **Model validation** | | |
|---|---|---|
| Clash score, all atoms | 10.25 ($97^{th}$ percentile) | 14.36 ($97^{th}$ percentile) |
| Poor rotamers | 0.22% | 0.06% |
| Favored rotamers | 99.11% | 98.77% |
| Ramachandran outliers | 0.16% | 0.10% |
| Ramachandran favored | 85.75% | 82.98% |
| MolProbity score | 2.18 ($100^{th}$ percentile) | 2.36 ($99^{th}$ percentile) |
| Cβ deviation | 0.00% | 0.00% |
| Bad bonds | 0.03% | 0.02% |
| Bad angles | 0.05% | 0.02% |
| Cis Prolines | 0.00% | 0.00% |
| Cis nonProlines | 0.01% | 0.01% |
| Twisted peptides | 0.09% | 0.19% |
| CaBLAM outliers | 5.30% | 5.6% |
| CA geometry outliers | 1.20% | 1.81% |
